# Recruitment, rewiring, and deep conservation in flowering plant gene regulation

**DOI:** 10.1101/2024.10.08.617089

**Authors:** Leo A Baumgart, Sharon I Greenblum, Abraham Morales-Cruz, Peng Wang, Yu Zhang, Lin Yang, Cindy Chen, David J Dilworth, Alexis C Garretson, Nicolas Grosjean, Guifen He, Emily Savage, Yuko Yoshinaga, Ian K Blaby, Chris G Daum, Ronan C O’Malley

## Abstract

Transcription factors (TFs) are proteins that bind DNA to control where and when genes are expressed. In plants, dozens of TF families interact with distinct sets of binding sites (TFBSs) that reflect each TF’s role in organismal function and species-specific adaptations. However, defining these roles and understanding broader patterns of regulatory evolution remains challenging, as predicted TFBSs may lack a clear impact on transcription, and experimentally-derived TF binding maps to date are modest in scale or restricted to model organisms. Here, we present a scalable TFBS assay that we leveraged to create an atlas of nearly 3,000 genome-wide binding site maps for 360 TFs in 10 species spanning 150 million years of flowering plant evolution. We find that TF orthologs from distant species retain nearly identical binding preferences, suggesting that regulatory evolution primarily arises from gain and loss of TFBSs. Within lineages however, conserved TFBSs are over-represented and found in regions harboring signatures of functional regulatory elements. Moreover, genes with conserved TFBSs showed a striking enrichment for cell type-specific expression in single-nuclei RNA atlases, providing a robust marker of each TF’s activity and developmental role. Finally, we compare distant lineages, illustrating how ancient regulatory modules were recruited and rewired to enable adaptations underlying the evolutionary success of grasses.

## Introduction

Evolution has produced an extraordinary array of functional diversity within the plant kingdom, from subtle distinctions between neighboring cells to large scale anatomical novelty. Underlying this variation is a dynamic system of gene expression orchestrated by transcription factors (TFs), proteins that promote or repress transcription by binding DNA at specific binding sites (TFBSs) containing a preferred sequence motif. Yet despite the vast morphological and physiological breadth of flowering plants, the same TF families regulate gene expression, raising the question of whether changes in TF binding preferences (trans evolution) or changes in TFBSs (cis evolution) serve as key drivers of plant diversification. Few large-scale efforts have tackled this question directly^1^, lacking the requisite TF binding maps from multiple plant lineages. In bacteria, a multiplexed DNA affinity purification (multiDAP) method^2^ successfully identified both conserved and lineage-specific TFBSs, building on DAP-seq^3^ by assaying the binding of vitro expressed TFs to genomic DNA fragments from multiple pooled species. Here, we apply the multiDAP method in 10 flowering plant genomes including both close relatives and distant lineages separated by 150 million years of evolution^4^, generating a broad resource of TF binding specificities and genome-wide TF binding maps. The resulting “pancistrome” allows us to dissect the relative contributions of trans and cis regulatory divergence within and between distant plant lineages.

Our cross-species TF binding maps further offer an unprecedented framework for comparative regulatory genomics. Defining core sets of conserved target genes for each TF, we track their expression across single-nuclei transcriptomes spanning multiple tissue types and developmental stages of five plant species, unravelling the regulatory programs controlling cell type identity across distantly related lineages. We further uncover clear patterns of TF network rewiring and recruitment to novel tissues, revealing how ancient and lineage-specific gene networks shape developmental programs and drive phenotypic adaptations.

## Results

### A plant pancistrome

In this study we optimized multiDAP for large eukaryotic genomes by refining experimental conditions to achieve low background and high sensitivity. We then applied it to investigate the function of 360 TFs representing 33 major TF families and 49 distinct motifs^3^, mapping their binding profiles across the genomes of 10 plant species simultaneously (**Supplementary Table 1**). This set of plants included *Arabidopsis thaliana* plus three other members of the Brassicaceae family (hereafter termed “brassica”): *Arabidopsis lyrata*, *Capsella rubella*, *Brassica oleracea* (wild cabbage), four additional members of the eudicots: *Solanum lycopersicum* (tomato), *Solanum tuberosum* (potato), *Fragaria vesca* (wild strawberry), *Populus trichocarpa* (black cottonwood), and two members of the monocots in the grass family Poaceae: *Oryza sativa* (rice) and *Sorghum bicolor* (sorghum) (**Supplementary Table 2**). To capture all TFBSs genome-wide, we used PCR-amplified genomic DNA fragments in the multiDAP assay to remove any tissue- or condition-specific DNA modifications such as DNA methylation that could impact TF binding. These comprehensive maps of TFBSs across 10 plant genomes allowed us to investigate drivers of regulatory evolution, assessing the relative contributions of trans and cis divergence.

### Minimal trans divergence

To measure the degree of trans evolution, we selected orthologous TFs from different species and examined divergence of their preferred motifs and genome-wide binding properties. We selected 35 *A. thaliana* TFs representing 21 different families and 26 distinct motifs, and identified the closest ortholog of each in five additional species, of which 108 performed well enough in duplicate multiDAP experiments to generate high quality binding maps: 21 from *A. lyrata*, 22 from *C. rubella*, 21 from *B. oleracea*, 21 from *S. lycopersicum*, and 23 from *O. sativa*. Across all orthologous TFs tested, we observed nearly identical motifs (**Fig. 1a**). The binding of *A. thaliana* TFs was highly predictive of the binding profiles of their orthologs in a given species’ genome (mean Pearson r=0.88, **Extended Data Fig. 1a and b**). This was true even between *A. thaliana* and *O. sativa* orthologs (r=0.85), which on average share only 52% amino acid identity. These correlations were nearly as high as between technical replicates of the same TF (r=0.92) and substantially higher than those measured between *A. thaliana* TFs within the same family (r=0.35), even when only considering TFs that also have highly similar consensus motifs (r=0.55, **Extended Data Fig. 1c**). This suggests that divergence of binding specificities between orthologous TFs plays a limited role in plant regulatory evolution.

**Fig 1.**
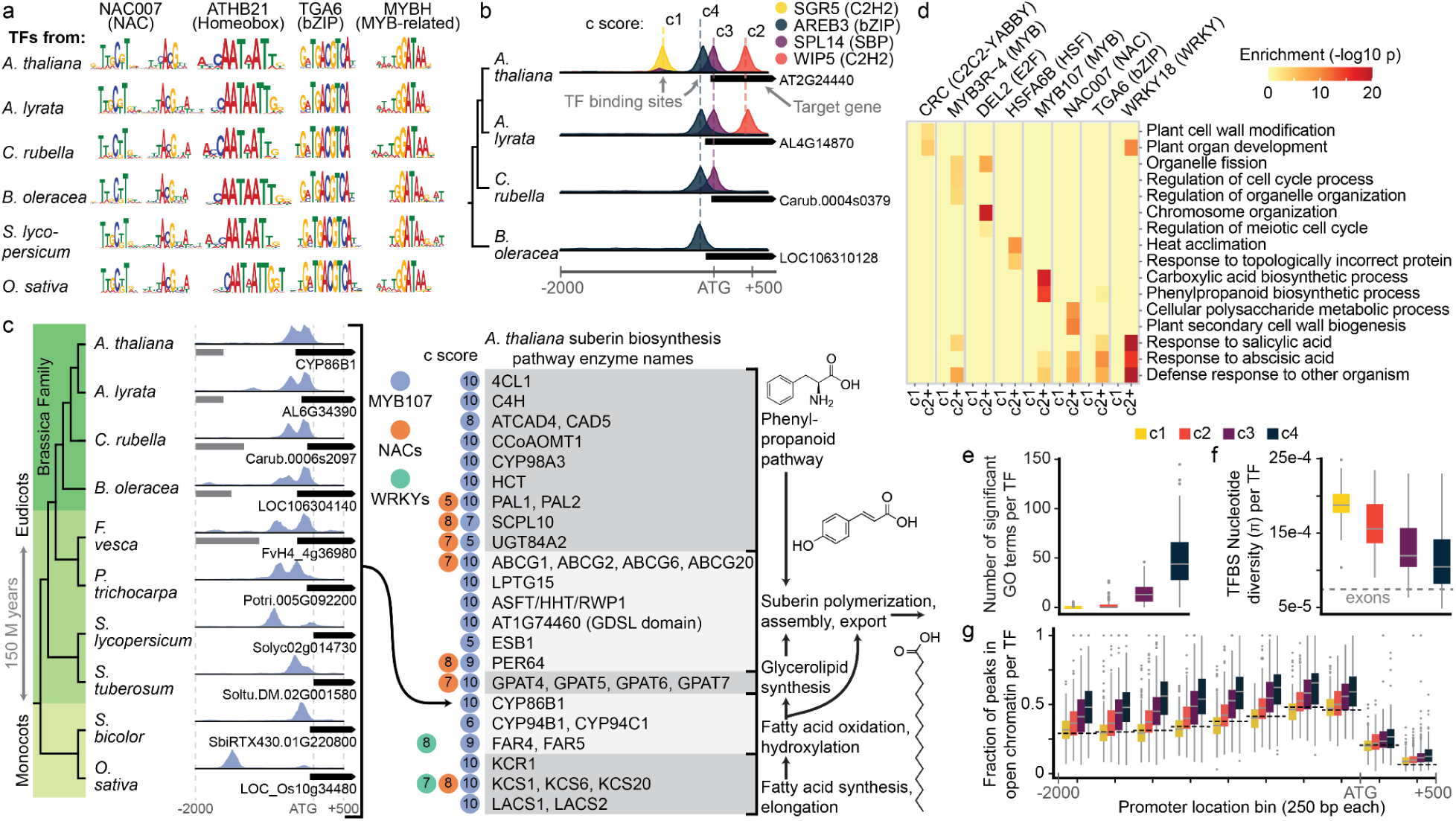
Plant multiDAP identifies motif specificities and TFBSs across 150 million years of evolution. See also **Extended Data Fig. 1** and 2 and **Supplementary Tables 1-4**. (a) High similarity of consensus motifs from four example sets of orthologous TFs from six species (b) Example of peak assignment to genes and conservation scores (c scores). (c) Left side shows phylogenetic relationships between 10 species used in this study. Track plots show as an example the target gene CYP86B1, which encodes an enzyme in the suberin pathway in *A. thaliana*, and its orthologs all targeted by MYB107. Right side shows key enzymes in suberin biosynthesis, which are ultra-conserved targets of MYB107 and other TFs in the NAC and WRKY families. Colors of circles and inset numbers next to each enzyme name indicate TFs and c scores, respectively. (d) GO term enrichment of non-conserved target genes (c1) versus conserved target genes (c2+). (e) Number of GO terms significantly enriched in target gene sets. Colors for each c score shown in legend are used throughout panels e-g. Boxplots in panels e-g show median +/− interquartile range with n = 244 TFs. (f) Average nucleotide diversity of TFBSs regions in the 1001 *A. thaliana* genomes. (g) Enrichment of open chromatin within TFBS regions, separated by c score and binned by promoter location. Dotted lines show expected frequencies of open chromatin overlap in each bin calculated from a random background distribution.

### Cis evolution across timescales

We next assessed TFBS conservation and divergence between closely related species and across diverse flowering plant lineages. The results above show that the binding of *A. thaliana* TFs in a different plant genome can serve as a strong predictor for the binding of the corresponding native TFs. Accordingly, we generated a more comprehensive multiDAP dataset of *A. thaliana* TFs covering the full range of families and distinct motifs. To dissect cis evolution at shorter timescales, we focused on the four closely related brassica species, using 244 *A. thaliana* TFs that performed well in our previous DAP-seq study^3^. We verified that multiplexing four genomic DNA samples did not significantly alter binding profiles or decrease signal (**Extended Data Fig. 2a**). To explore the opposite extreme of regulatory conservation, we measured the binding of 74 representative *A. thaliana* TFs in the genomes of all 10 plants. This subset was selected to include well-studied TFs that showed distinct binding profiles in *A. thaliana*, and while not all have a direct ortholog in all 10 profiled species, each represents the set of native TFs with the same motif preference. Although running the multiDAP assay with 10 genomes simultaneously introduced a slight decrease in signal for some TFs, overall binding profiles remained similar (r=0.87, **Extended Data Fig. 2a**).

We detected TFBSs throughout all genomes, including in genes, promoters, and distal intergenic sites upstream and downstream of genes. While the total number of TFBSs per genome was highly variable and dependent on genome size, the density of TFBSs near gene starts was more stable (**Extended Data Fig. 2b**). To quantify cis evolution at both short and long time scales, we first developed a reliable approach to identify TFBSs shared across species. We focused on TFBSs near the start of protein-coding genes, from 2,000 bp upstream to 500 bp downstream of protein start codons. We assigned all protein-coding genes from all species to orthogroups (an assignment based on shared amino acid sequences)^5^. Then, for each *A. thaliana* TF-target gene relationship we generated a conservation score (c score), representing the number of species in which an orthologous gene had a binding site for the same TF (**Fig. 1b**). Target genes scored as c1 represent genes where a binding site for a given TF could only be found in *A. thaliana*, whereas target genes scored as c4 represent genes where binding sites for a given TF were found in *A. thaliana* plus orthologous genes from three other species. Our approach allowed us to detect conservation even across highly diverged species where intergenic sequence alignments cannot reliably identify conserved TFBSs^6–8^.

Using our brassica atlas, we assigned c scores to 1.1 million total TF-orthogroup interactions represented in the *A. thaliana* genome. Of these, 24% were conserved across all four brassica genomes (c4), with the next largest category at 21% that were private to *A. thaliana* (c1) (**Extended Data Fig. 2c**). By comparing the number of TF-target genes shared between two or more (c2+), three or more (c3+) or all four species (c4) to a background frequency calculated from overlap between sets of randomly generated TF target orthogroups, we found that all conserved TF target gene sets were significantly larger than expected by random chance (p < 0.01). Furthermore, the enrichment and significance increased as the c score increased (**Extended Data Fig. 2d**). This indicates that while considerable cis evolution has occurred since these brassica diverged 21 million years ago^9^, a large set of core TF target genes were retained.

The 150 million year evolutionary distance between the monocots and eudicots^4^ within our flowering plant multiDAP atlas enabled identification of smaller ultra-conserved TF target gene sets. Among the *A. thaliana* target orthogroups conserved in the four tested brassica species, 7.1% were further conserved across all 10 species (c10 target genes). Many are consistent with previously elucidated TF functions and capture ancient pathways shared across flowering plants. For example, MYB family TF MYB107, along with several NAC and WRKY family TFs, consistently target a core set of genes associated with suberin production^10–12^, including key enzymes at all major steps in the suberin biosynthesis pathway, the upstream phenylpropanoid and fatty acid synthesis pathways, as well as transporters involved in exporting suberin monomers across the cell membrane (**Fig. 1c**)^10,13,14^. We further uncovered many ultra-conserved yet previously unknown MYB107 target genes with predicted suberin-related functions, which may serve as attractive targets for future characterization. Other TFs that have retained a core set of ultra-conserved target genes play key roles in coordinating development or environmental responses, such as PC-MYB1/MYB3R1 in cell division^15^, GTL1 in root hair development^16^, Dehydration Response Element 1A (DREB1A) in drought response^17^, and CAMTA1/EICBP.B in stress hormone responses^18^. Overall, while TFBSs have diverged more rapidly than TF binding specificities, our atlas revealed substantial conservation of TF target genes in closely related species, along with a smaller set of ultra-conserved core regulons that have persisted across 150 million years of flowering plant evolution.

### Conservation highlights functional TFBSs

We hypothesized that TFBSs regulating critical gene functions would be among those conserved between closely related species, reflecting the selective pressure to maintain key regulatory interactions, while TFBSs with little impact on gene expression would not be subject to the same selective pressure. To investigate this, we applied several independent approaches to test whether TFBSs conserved within the brassica dataset exhibited signatures of functional cis-regulatory elements (CREs). First, we found that while c1 target gene sets were rarely enriched for functional gene ontology (GO) terms, conserved *A. thaliana* target gene sets with c scores of c2 or greater were significantly enriched for GO terms that are consistent with known TF functions, suggesting that conservation across even two closely related species highlight functional binding sites (**Fig. 1d**, see also **Supplementary Table 3**). Among these were TFs spanning key developmental and environmental response pathways, with GO terms related to organ development for CRC^19^, cell division for MYB3R-4^20^ and DEL2^21^, heat stress for HSFA6B^22^, suberin synthesis for MYB107^10^, and response to pathogens and stress related hormones for TGA6^23^, WRKY18^24^ and NAC007/VND4^25^. The number of significantly enriched GO terms increased as the conservation level increased (**Fig. 1e**). Some of these functional associations extend past brassica, and were strongly enriched among the ultra-conserved c10 target genes in the 10 species dataset (**Supplementary Table 4**). Second, we found evidence of purifying selection acting to preserve the sequences immediately underlying conserved peak summits across wild *A. thaliana* populations^26^. We observed decreasing nucleotide diversity at peaks with higher c scores, with significantly lower nucleotide diversity underlying c4 TFBSs compared to all other TFBS sets (t-test, p < 0.05), nearly as low as that found within protein-coding exonic regions (**Fig. 1f**). Finally, conserved binding sites for most TFs showed enrichment in accessible chromatin regions (ACRs) identified by ATAC-seq^27^. Since both TFBSs and ACRs are known to be concentrated near transcription start sites^27^, we split *A. thaliana* promoter regions into 250 bp bins and found that conserved TFBSs in all promoter bins were enriched in ACRs, while c1 TFBSs were not (**Fig. 1g**). Taken together, this demonstrates that conserved TFBSs are enriched for CREs and highlights the power of using conservation to identify functionally important TF target genes.

### Charting TF activity in single nuclei

As conserved TFBSs are enriched for the hallmarks of functional CREs, TF binding at these sites is likely to impact expression of the associated target gene. We therefore explored whether quantifying the expression of each TF’s conserved target genes would allow us to chart TF activity across cell and tissue types in multiple species. To do so, we generated snRNA-seq atlases for seedling root and shoot, mature leaf, and flower bud tissues of all four brassica species, profiling in total 163,982 high-quality single nuclei transcriptomes, with a median of 899 unique transcripts and 742 genes per nucleus (**Supplementary Table 5**). Using published reference atlases^28–30^ and marker gene lists^31^, we assigned all *A. thaliana* nuclei to 61 unique cell type labels (**Fig. 2a, Extended Data Fig. 3a-c**), including a small number of seedling labels that were not present in published protoplast-based atlases, but were supported by integration of an external seedling root nuclei dataset^32^. We used SAMap^33^ to lift-over labels from the *A. thaliana* atlases to the other three species (**Fig. 2b**), resulting in 57 labels with at least 10 nuclei from all four species (**Supplementary Table 6**).

**Fig. 2.**
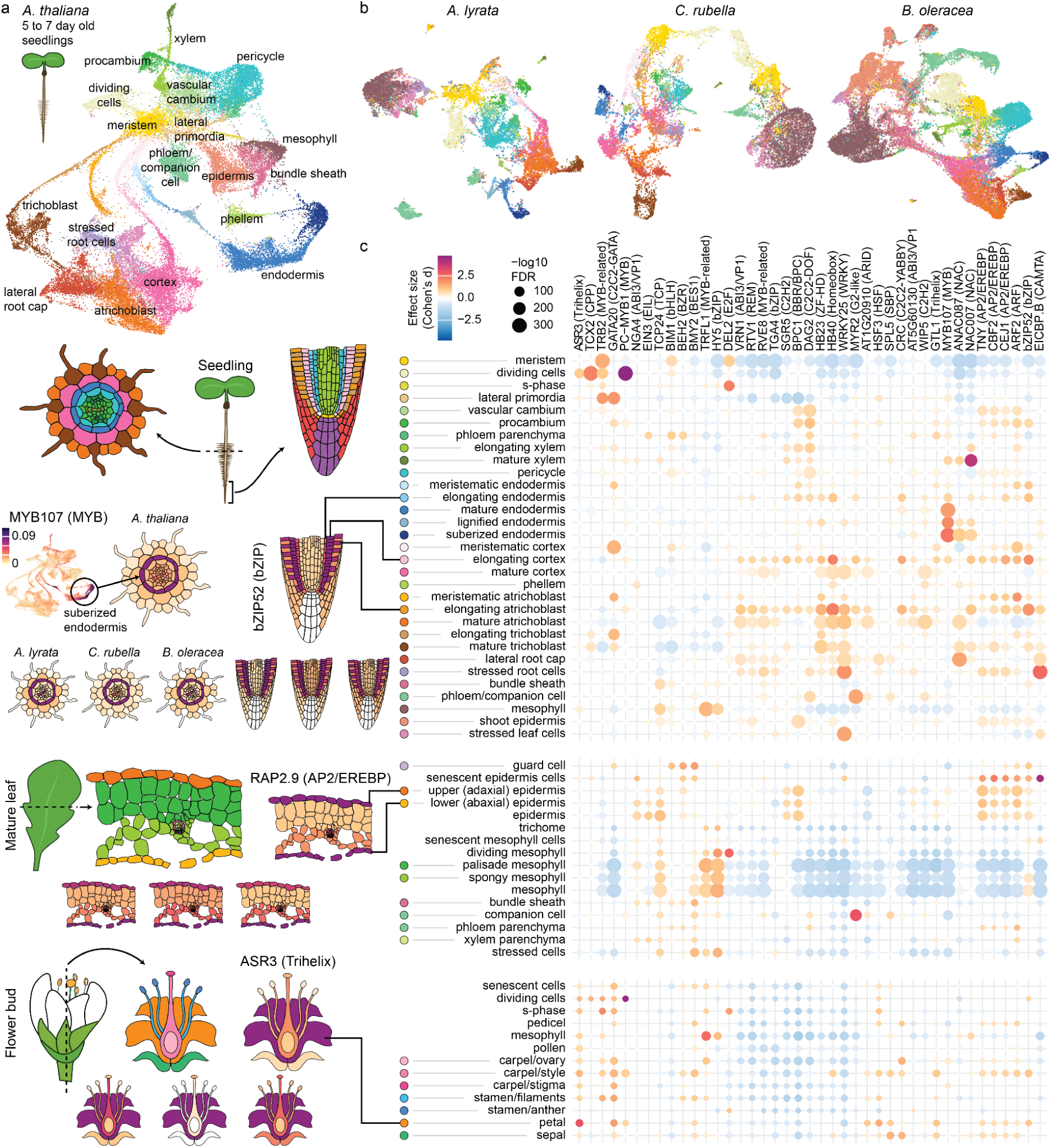
Expression of TF target genes across snRNA-seq seedling atlases enable construction of a multi-tissue regulatory map of cell type-specific gene expression programs. See also **Extended Data Fig. 3-5** and **Supplementary Tables 5-7**. (a) UMAP representation of the transcriptomes of >45,000 single *A. thaliana* seedling nuclei corresponding to 32 labeled cell types and (b) UMAP representations of corresponding data from three brassica relatives, with cell type labels transferred from the *A. thaliana* atlas. Colors in panels a and b match the colored circles shown beside cell type labels in the seedling portion of the heatmap below. (c) Heatmap on right shows enrichment or depletion of c4 target gene expression for representative TFs (horizontal axis) across labeled cell types in *A. thaliana* seedling, leaf, and flower bud snRNA-seq atlases (vertical axis). Dot size represents the FDR-corrected −log10 p-value from a Wilcoxon rank sum test of summarized target expression scores for each cell type label vs. all other cells, and color represents the effect size of the expression enrichment (warm colors) or depletion (cool colors). Enrichments with FDR-corrected p-value < 0.01 are shown. Left half shows detailed plots for one example TF in each of four different anatomical regions. The spatial arrangement of cell types in a cross section of mature root is shown in a graphic at the top far left, with cell types colored according to circles beside cell type labels in the vertical axis of the corresponding seedling portion of the heatmap. Summarized expression of TF MYB107 c4 target genes in each cell is shown in the *A. thaliana* UMAP plot directly below. Strong enrichment of target expression in suberized endodermis cells is highlighted with a labeled circle. Average target gene expression scores for each cell type are then projected onto the spatial map of mature root cell types, shown directly to the right of the UMAP. Projections for the other brassica are shown directly below. The subsequent examples use the same structure, but highlight bZIP52 target genes in root tip, RAP2.9 target genes in mature leaf, and ASR3 target genes in flower bud.

We first confirmed that the expression profiles of conserved target genes generally reflected regulation by the associated TF. Within the *A. thaliana* seedling atlas (the *A. thaliana* tissue with the deepest per-cell transcript coverage), co-expression of TF-target gene pairs was significantly higher than background and strongest among the most conserved (c4) target genes, regardless of TF binding affinity (**Extended Data Fig. 4a**). Furthermore, in every species, c4 target gene sets showed expression patterns that were significantly more cell type-specific than c1, c2, or c3 target genes (**Extended Data Fig. 4b**), as would be expected from genes subject to strong transcriptional control by distinct TFs.

As a measure of TF activity, we then quantified the enrichment or depletion of expression of each TF’s c4 target genes in each cell type (**Supplementary Table 7**). The resulting set of TF activity scores yield an expansive map of TF regulatory roles in cell growth, differentiation, and function, with 241 of 244 tested TFs demonstrating significant activity in at least one cell type. While many associations reflect known regulatory biology (**Fig. 2c**), such as the role of WRKY TFs in stress response and senescence^24^, G2-like TF MYR2 in seedling and leaf phloem companion cells^34^, and PC-MYB1/MYB3R1 and TCX2 in cell division^35,36^, numerous previously uncharacterized TF roles were identified as well. For example, mesophyll cells at all developmental stages in every tissue (cotyledon, leaf, and flower) were strongly and specifically associated with both HY5 (bZIP) and TRFL1 (Myb-related). While HY5 is known to be active in mesophyll cells^37^, a role for TRFL1 or any close paralogs in mesophyll cells has not been previously documented. Extending our analysis to examine ultra-conserved TF target sets suggested that certain core regulons represent ancient cell type-specific processes, for example c9 target genes of secondary cell wall biosynthesis regulator NAC007/VND4^38^ were more specifically expressed in mature xylem cells than TF target sets at any other conservation level (**Extended Data Fig. 4c**). Accordingly, a TF’s conserved target genes often retained the same cell type specificity across multiple species’ atlases (**Fig. 2c**). For example, MYB107 target genes were strong and specific markers of late-stage endodermis cells (including lignified and suberized endodermis) in the seedling atlases of all four species. As several MYB family TFs are known to contribute to suberin deposition^11^, untested family members with similar binding specificity could be responsible for activating target gene expression. However, MYB107 expression was detected specifically in endodermis cells in multiple species’ seedling atlases, supporting a previously unrecognized role for this transcription factor beyond the seed coat^10^. bZIP52 target genes were specifically expressed in elongating endodermis, cortex, and atrichoblast in all four species, consistent with recent work demonstrating the role of Group I bZIP TFs in elongation of multiple root cell types^39^. Similarly, AP2/EREBP TF RAP2.9 target gene expression marked epidermal leaf cells in all four species reflecting a potential role in leaf cuticular wax biosynthesis^40^, and the target genes of ASR3 (Trihelix) showed petal-specific expression in all four species, consistent with Trihelix regulation of petal initiation^41^. In total, we identified key regulators of cell type-specific expression (FDR<.01, Cohen’s d >= 0.8) in 57 of 61 cell types, with all but 5 showing significant activity from more than one TF family. These results underscore the extensive coverage of our TFBS atlas and open new avenues for engineering targeted cellular responses in plants.

### Regulatory networks define cell types

Our map of conserved TF target gene expression enrichments suggests that each cell type has a unique fingerprint of TF activity, often involving multiple TFs acting together. We next asked whether TFs with enriched target gene expression in the same cell type operate through discrete or overlapping regulons. We visualized each cell type’s regulatory network, connecting enriched TFs to c4 target genes among the cell type’s top 100 marker genes. In some cell types, a single TF family appeared to be acting as a master regulator. For example, in dividing cells, 77 marker genes were targets of MYB3R5 (**Extended Data Fig. 5a**), a 15-fold enrichment over the expected background. Of the remaining 23 marker genes, at most five were targeted by any other single tested TF, suggesting that MYB3R5 and TFs with similar binding specificity are the primary drivers of gene expression within the cell division pathway. We hypothesized that regulation of such an ancient and ubiquitous pathway would be tightly conserved beyond brassica, and indeed, close to half (48%) of the markers targeted by MYB3R5 in *A. thaliana* had orthologs targeted by MYB3R5 in all 10 profiled plants. In most cell types, however, the strongest marker genes were frequently co-targeted by multiple TF families. For example, in procambium, 64 of the top 100 marker genes were targeted by one or more TFs from the BBR/BPC and C2C2-DOF TF families and 25 of these were co-targeted by both **(Extended Data Fig. 5b)**, significantly more than expected by chance given the number of markers targeted by each individually (p=6×10^−5^). Similarly, co-targets of both NAC and MYB family TFs were enriched among suberized-endodermis markers (p=0.007) (**Extended Data Fig. 5c**), and co-targets of both HOMEOBOX and MYB-RELATED family TFs were enriched among mature-atrichoblast markers (p=8×10^−7^) **(Extended Data Fig. 5d)**. Across the 23 seedling cell types with multiple enriched TF families, 706 (30.7%) of the top marker genes were co-targeted by TFs from more than one enriched TF family. These results suggest that cell type specific regulatory networks are deeply intertwined, with multiple non-redundant TFs acting both independently and in concert to regulate gene expression.

We next tested how well the complete set of all measured TF activity scores characterizes a cell type, regardless of species, by calculating correlations between pairs of TF activity score profiles from cell types in different brassica species. We found the highest correlations between matching cell types in different species **(Extended Data Fig. 5e**), indicating that core regulatory architecture of developmental programs has remained largely stable across the 21 million years separating these species, with the same TF families active in the same cell types across brassica. These findings confirm that TF activity score profiles capture the salient features of cell type-specific regulation.

### Recruitment and rewiring of TF networks

Having established that TFBSs conserved within the brassica family underlie regulatory networks that drive cell identity, we next investigated the fate of these conserved sites across the 150 million years separating brassica from monocots. We used our 10-species multiDAP dataset to assign grass-c2 scores to TFBSs shared by two model monocots in the grass family, rice and sorghum, and looked for overlap with TFBSs conserved in all four brassica species (brassica-c4 TFBSs) as well as lineage-specific TFBSs that we ascribe to TF rewiring (**Fig. 3a**). Counting only unique orthogroups targeted by a TF, we found that 9,027 (21%) brassica-c4 target orthogroups were conserved in both grasses, constituting a core regulon of target genes controlled by the same TF in all six species. An additional 50% of brassica-c4 target orthogroups were absent in both grasses, with half (10,530) corresponding to orthogroups lacking grass genes, while the majority of the rest represent shared orthogroups with lineage-specific TFBS loss or gain. Across the TFs, we observed a wide range of cross-lineage conservation, with anywhere from 4% to 80% of brassica-c4 target orthogroups conserved in both grasses, yet the vast majority (87%) had core regulons significantly larger than expected by chance (FDR<0.05, **Extended Data Fig. 6**). Overall, the widespread enrichment of cross-lineage TFBS conservation suggests that the majority of TFs control core regulons that persist alongside substantial rewiring.

**Fig. 3.**
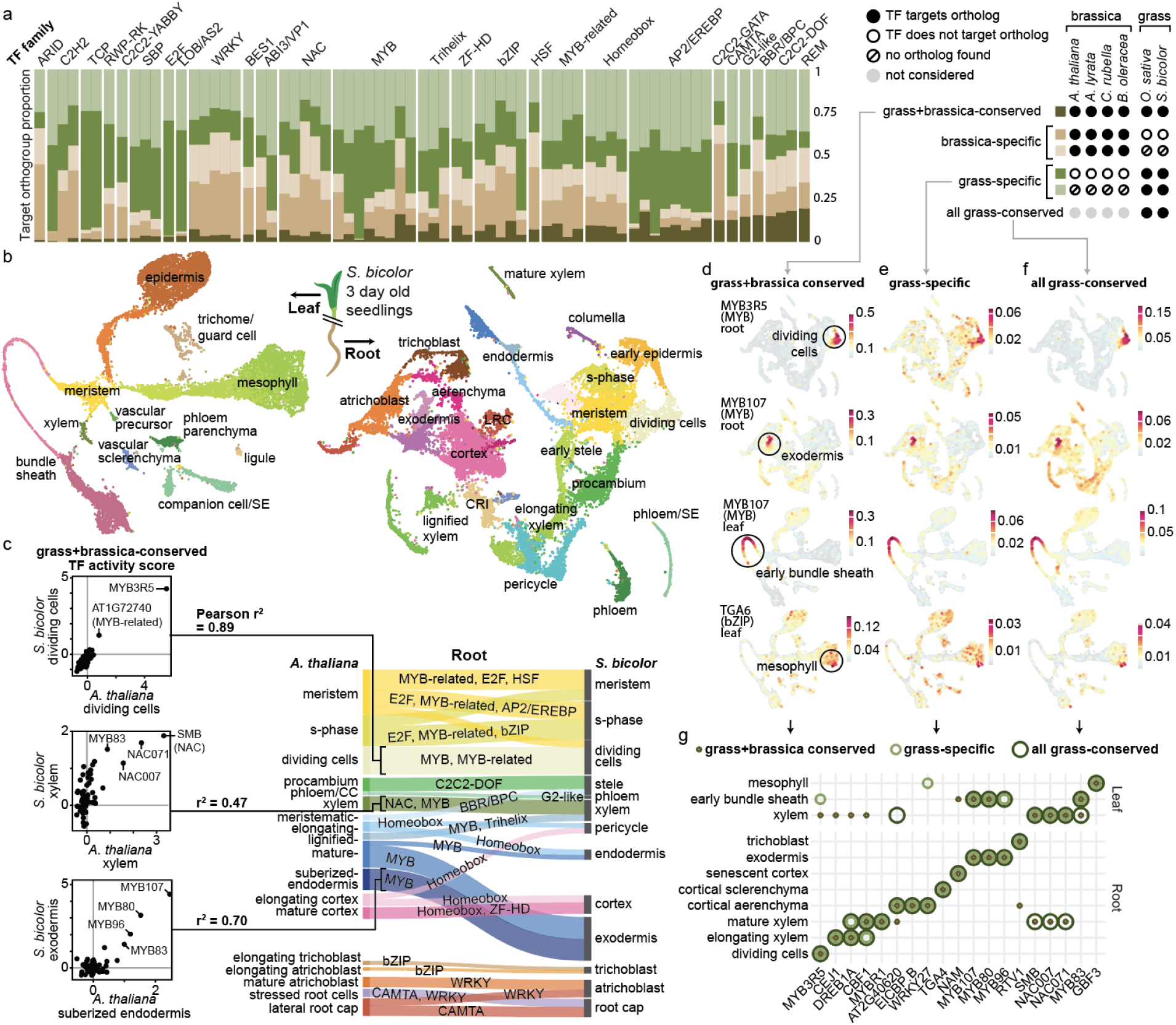
Single nuclei transcriptomes of *S. bicolor* seedlings reveal conserved and grass-specific regulatory networks driving cell type identity. See also **Extended Data Fig. 6-9**, **Extended Data Table 1**, and **Supplementary Table 9**. (a) Relative proportions of target orthogroups conserved within and between grass and brassica lineages, for each TF (horizontal axis). (b) UMAP representation of the transcriptomes of >32k single sorghum nuclei corresponding to 43 labeled cell types. SE = sieve element; LRC = lateral root cap; CRI = crown root initials (c) Correlations of TF activity scores calculated for sorghum and *A. thaliana* cell types using target genes conserved in both grass and brassica lineages. Left: three examples are shown as scatterplots, where each dot represents the activity score (measured by Cohen’s d) of a TF in an *A. thaliana* cell type (x-axis) vs. a sorghum cell type (y-axis). Right: Diagram linking root cell types between *A. thaliana* and sorghum, where the widths of the connections correspond to the TF activity score profile correlations (Pearson r^2^) between each pair. Only correlations with r>0.3 are shown, after filtering to the top two correlations per sorghum cell type. (d) Cross-lineage conserved target genes identified by multiDAP using *A. thaliana* TFs display cell type-specific expression patterns in sorghum. (e) Expression patterns of grass-specific TF target gene sets for the same TFs shown in panel c. (f) Expression patterns of all TF target genes conserved in both grasses for the same TFs shown in panels c and e. (g) Examples where strong enrichments for all three TF target gene subsets appear in the same sorghum cell type. Only enrichments with FDR<0.05 and Cohen’s d > 0.5 are shown, after filtering to the top two enrichments per TF across all root and leaf cell types.

To explore the functional consequences of conservation and rewiring, we generated snRNA-seq datasets from leaves and roots of 3 day old sorghum seedlings, comprising a total of 13k and 19k nuclei, respectively, after stringent quality filtering and identified 43 cell types (**Fig. 3b, Extended Data Fig. 7, Supplementary Table 8**). We first asked whether the same TFs were active in brassica and grass cell types, using core regulon target genes as a marker of TF activity. Comparing activity scores across 74 TFs for each *A. thaliana* and sorghum cell type (**Supplementary Table 9**), we found the highest correlations between many corresponding cell types (**Fig. 3c**, **Extended Data Fig. 8**). These correlations were primarily driven by small collections of TF families controlling different cell layers, including MYB, MYB-related, and E2F in meristematic and immature cell types, C2C2-DOF, G2-like, and NAC in vascular layers, and WRKY, bZIP, and CAMTA in terminal developmental stages of the epidermis and root cap. Furthermore, many core regulons for individual TFs showed strong cell type-specific expression in sorghum. For example, expression of MYB3R5 target genes was enriched in sorghum dividing cells, while MYB107 target genes displayed strong expression in bundle sheath and exodermal layers, and TGA6 target genes marked a discrete set of cells within the mesophyll (**Fig. 3d**).

We also observed several striking examples of regulatory rewiring reflecting distinct patterns of cell type evolution. For example, MYB83 and NAC007 TFs are active and strongly expressed in mature xylem cells of both species (**Extended Data Fig. 9**), consistent with the known roles of NAC and MYB families in secondary wall biosynthesis^42,43^. However, our data revealed that the dominant family shaping xylem identity is reversed in the two plant lineages **(Extended Data Table 1)**. While NAC007 and related factors play the primary role in *A. thaliana* (with 27 of the top 100 xylem marker genes carrying a brassica-c4 NAC007 TFBS, compared to 4 with a MYB83 TFBS), MYB83 and related factors appear to dominate in sorghum xylem (where only seven of the top 100 xylem markers have a grass-c2 NAC007 TFBS, yet 14 have a grass-c2 MYB83 TFBS). Moreover, we find ample evidence of direct TFBS switching within individual pairs of orthologous genes. Among the top 50 *A. thaliana* xylem markers, we identified seven cases in which the *A. thaliana* gene is a target of NAC007 but not MYB83, while the sorghum ortholog is a target of MYB83 but not NAC007, an 81-fold enrichment compared to all *A. thaliana* genes with a sorghum ortholog. Six of these seven sorghum orthologs, including IRX6, IRX3, LAC17 and NAC073, were still strong markers for xylem in sorghum. In fact, NAC073 has two sorghum orthologs: one ortholog (SbiRTX430.09G246200) mirrors the *A. thaliana* gene with a NAC007 TFBS and no MYB83 TFBSs and is moderately enriched in xylem (log2fold-enrichment=1.8). The second (SbiRTX430.03G271900) shows the reverse TFBS profile (no NAC007 TFBS and a grass-specific MYB83 TFBS) and is more than twice as enriched in xylem (log2fold-enrichment=4.1). Importantly, these lineage-specific differences cannot be attributed to technical artifacts, as with the multiDAP approach, all species are jointly incubated with each TF and sequenced as a pool.

A second example revealed recruitment of developmental pathways underpinning grasses’ ability to survive in hot, dry conditions. Drought tolerance is in part enabled by an exodermal cell layer that produces suberin, a waxy-substance that regulates water and nutrient transport and provides a protective barrier to roots^44^. While this layer is absent in *A. thaliana*, it instead suberizes an internal cell layer, the endodermis. Consistent with this, we observed a strong correlation of TF activities between sorghum exodermis and *A. thaliana* endodermis (see **Fig. 3c**), primarily driven by MYB family TFs which regulate suberin and lignin production^45^. Furthermore, we found evidence that this MYB-controlled network has been recruited to the bundle sheath of sorghum leaves (see **Fig. 3d**), where suberin forms the gas-tight layers that support the more efficient C_4_ carbon fixation pathway employed by many grasses^46^. Recruitment of the MYB107 regulon was accompanied by substantial regulatory rewiring. 31 of the top 100 sorghum exodermis markers and 24 of the top 100 early bundle sheath markers were associated with MYB107 TFBSs that were detected in both grasses but none of the brassica species. In both cases these grass-specific target genes outnumbered MYB107 core regulon genes among the top cell type markers. While the majority lacked brassica orthologs altogether, in both cell types the subset with known brassica orthologs showed clear evidence of regulatory divergence, as none of these orthologs were strongly expressed in A. thaliana endodermis.

In fact, we found a large number of grass-specific TFBSs for almost every TF. If these represent true TF target genes, we reasoned that their expression should generally mirror that of core regulon target genes. In addition to MYB107, several TFs including MYB3R5 and TGA6 showed very clear alignment between the expression patterns of grass-specific and core regulon target genes (**Fig. 3e and f**). A more comprehensive search revealed 17 additional examples of TFs where the strongest enrichments aligned in the same cell type (**Fig. 3g**). In some cases, these grass-specific target genes outnumber core regulon genes by a factor of 10:1, yet strong evidence of shared transcriptional control was observed. Together, this suggests that many grass-specific TFBSs reflect functional CREs that place new target genes under the control of TFs driving core developmental programs. Importantly, these results hinge on the ability of *A. thaliana* TFs to identify grass-specific target genes that may not even exist in any brassica genome.

Collectively, these findings reveal how selective repurposing of ancient gene networks supports the emergence of new cell types and functions, which together with widespread cis-regulatory rewiring give rise to lineage-specific traits. In conjunction with complementary studies in systems spanning photosynthetic cell types^47^ to human neurons^48^, this supports an emerging model of how ancestral TFs are harnessed during the evolution of specialized phenotypes.

## Discussion

Our study presents a comprehensive account of how TF regulatory networks shape cell type identity and evolution, leveraging new binding maps for 360 TFs and 14 new single nuclei transcriptomic atlases across multiple tissues and both major lineages of flowering plants. Together, these provide a foundational resource for unraveling regulation through the lens of comparative genomics. By integrating these atlases, we demonstrated that conserved target genes are powerful predictors of TF pathway functions, and that core TF regulons persist alongside ample rewiring throughout the course of plant evolution.

Further work is required to fully understand the complexities of gene regulation. While we chose to focus on promoter regions near protein-coding genes, our plant multiDAP atlas includes distal binding sites which may also contribute to gene regulation. Secondly, overlapping binding specificities of multiple TFs within a family complicates the definitive attribution of regulatory outputs to specific TFs. Nevertheless, we observed many cases where conserved binding sites for specific TFs known to play roles in a developmental response program are highly predictive of relevant gene expression patterns even in divergent species. Finally, while we focused here on conserved TFBSs, both our previous bacterial multiDAP study^2^ and a DAP-seq study of two maize strains^49^ identified strain-specific binding sites that impact expression and phenotype. Future studies combining our atlas with genetic, epigenetic, and functional resources may help to identify patterns associated with novel functional binding sites. Expanding this atlas to include more species, developmental stages, and environmental conditions will deepen our understanding of how regulatory network evolution generates phenotypic diversity, with broad implications for synthetic biology and agricultural biotechnology.

## Acknowledgments

The work conducted by the U.S. Department of Energy Joint Genome Institute (https://ror.org/04xm1d337), a DOE Office of Science User Facility, is supported by the Office of Science of the U.S. Department of Energy operated under Contract No. DE-AC02-05CH11231.

We also thank the following individuals: Chris Beecroft and Tatiparthi Reddy for assisting with submission of raw data files and curation of associated metadata; Laura Gerard for illustrating plant cartoons; Benjamin Cole, Jacqueline Humphries, Axel Visel, and Zhuzhu Zhang for editing the text and providing comments; Missy Fix, Nahla Bassil, and Kim Hummer at the USDA ARS NCGR for providing *F. vesca* germplasm; Joseph Edwards and the Sundar lab at UC Davis for providing *O. sativa* germplasm; Jesse Schartner at the USDA ARS VCRU for providing *S. tuberosum* germplasm; Melanie Harrison and Tiffany Fields at the USDA ARS PGRCU for providing *S. bicolor* germplasm.

## Author contributions

Conceptualization LAB, RCO, SIG; Methodology AMC, DD, LAB, PW, RCO, SIG, YZ; Validation AMC, DD, LAB, RCO, SIG, YZ; Formal Analysis AG, AMC, LAB, LY, RCO, SIG; Investigation CC, ES, GH, NG, PW, YZ; Writing, AMC, LAB, PW, RCO, SIG, YZ; Visualization, AMC, LAB, LY, SIG; Supervision CGD, IB, LAB, RCO, SIG, YY

## Data and Code Availability

Scripts and example data files used for DAP-seq and single nuclei analyses are available in a git repository at https://code.jgi.doe.gov/LBaumgart/plant-multidap-and-single-cell.

Raw DAP-seq fastq sequence data files were submitted to the National Center for Biotechnology Information under BioProject no. <u>PRJNA1177505</u>, for which curated metadata are managed in GOLD (https://gold.jgi.doe.gov/).

Single-nuclei datasets are in the process of submission to NCBI and will be available at the time of publication. In the meantime, they can be accessed at: https://portal.nersc.gov/cfs/m342/leo/plant_multidap_and_single_cell/

Data tables containing filtered and annotated peaks assigned to genes with c scores from the 4 species and 10 species datasets, along with raw narrowPeak files are being submitted to Zenodo at <DOI TBD>. In the meantime they are available at: https://portal.nersc.gov/cfs/m342/leo/plant_multidap_and_single_cell/

## Methods

### Plant Germplasm and Reference Genomes

Reference genome sequences and annotations for *A. thaliana*^50^, *A. lyrata*^51,52^, *C. rubella*^53^, *B. oleracea*^54^, *F. vesca*^55^, *P. trichocarpa*^56^, *S. lycopersicum*^57^, *S. tuberosum*^58^, *S. bicolor* (These sequence data were produced by the US Department of Energy Joint Genome Institute), and *O. sativa*^59^ were downloaded from Phytozome or NCBI RefSeq. Gerplasm was obtained either from ABRC, USDA, TGRC, or inhouse collections. See **Supplementary Table 2** for a full list of germplasm and reference genome sources.

### DAP-seq Experiments

The DAP-seq and multiDAP-seq assays were performed using the reagents as previously described^2,3,60^, but with modifications to optimize sensitivity with all 96-well plate operations taking place on a Hamilton Vantage liquid handler. Genomic DNA (gDNA) was extracted from each of the 10 plant species using a CTAB protocol, after which gDNA fragment libraries were constructed with an average shearing size at 150 bp. Different species were labeled by unique Illumina i5 index adapters. Upstream of the DAP-seq assay, fragment libraries were amplified for 10 cycles to remove any DNA modifications.

The coding sequence (CDS) of plant transcription factors were cloned into the pIX-HALO-PaqCI vector, a modified vector optimized for chewback cloning but resulting in identical assembled plasmid sequences as the original pIX-HALO^3,60^. The HaloTag-fused TF CDSs were amplified by the primers pIX-Halo-T7-fwd (5’-GTGAATTGTAATACGACTCACTATAGGG) and pIX-Halo-AfterPolyA-rev (5’-CAAGGGGTTATGCTAGTTATTGCTC) using KAPA HiFi HotStart ReadyMix (Roche KK2602). Resulting PCR products were cleaned up using the Mag-Bind TotalPure NGS Kit (Omega M1378-02), and correct amplicon sizes were verified on a Fragment Analyzer system (Agilent) using the HS NGS Fragment Kit (Agilent DNF-474-1000). A total of 10 µL at a minimum concentration of approximately 100 ng/µL of each PCR amplicon was used to express TF proteins in vitro using the TnT T7 Quick for PCR DNA System (Promega L5540) with an incubation at 30°C for 2 hours. Each 96 well plate of protein included four negative control wells containing a mock in vitro protein expression without any DNA expression template, to be used for background signal subtraction in the downstream peak-calling step.

Magne HaloTag Beads (Promega G7282) were first washed and resuspended in 50 µL of binding/wash buffer (1x phosphate buffered saline + 0.005% NP40 detergent). To bind TFs to magnetic beads, 20 µL of washed beads were incubated with 100 µL of each protein expression reaction at room temperature for 1 hour on a rotator. The HaloTag-TF-bound beads were gently washed in 150 µL of the same binding/wash buffer 3x, using a magnetic plate adapter to pellet the beads and discard the wash buffer between each wash. The beads were then combined with the gDNA libraries in a volume of 50 µL, brought to a total volume of 150 µL using binding/wash buffer, and incubated on a rotator at room temperature for 1 hour. Input gDNA library amounts were titrated based on genome size to target equal sequence coverage of each species: *A. thaliana* 100 ng, *A. lyrata* 172 ng, *B. oleracea* 407 ng, *C. rubella* 112 ng, *F. vesca* 183 ng, *O. sativa* 312 ng, *P. trichocarpa* 327 ng, *S. lycopersicum* 652 ng, *S. tuberosum* 618 ng, *S. bicolor* 565 ng. In addition, 10 µg of salmon sperm DNA (Invitrogen 15632011) was added alongside gDNA libraries to reduce background signal caused by non-specific binding. After incubation, the beads were again gently washed with the same wash buffer 3x, after which the wash buffer was removed and discarded. We found that the wash conditions are critical to achieving high sensitivity. At each wash step, the bead pellet should be fully resuspended by gentle pipetting of the wash buffer stream directly at the bead pellet. After the final wash, the beads were resuspended and boiled in 22 µL of i7 index primers^2^ at 95°C for 15 min, and 20 µL of eluate from each well was transferred to a new plate and mixed with an equal volume of KAPA HiFi HotStart ReadyMix for 10 cycles of amplification. Equal volumes of each reaction were pooled across rows of the plate, and then purified by gel purification. Pools were sequenced either on an Illumina NovaSeq 6000 S4 flow cell or NovaSeq X plus 10B or 25B flow cell, using 2×150 bp paired-end sequencing, targeting 10 million reads (5 million fragments) per TF for *A. thaliana*, and proportionally scaled to larger genomes as dictated by the relative gDNA library input mass. For the four species brassica dataset, we identified libraries from that first round of sequencing with low sequencing yield and selectively pooled these for a second round of sequencing to increase coverage.

### Single nuclei Experiments

#### Plant growth conditions

*A. thaliana*, *A. lyrata*, *C. rubella*, and *B. oleracea* seeds were surface sterilized before planting vertically on agar plates containing 3 mM Ca(NO_3_)_2_, 1.5 mM MgSO_4_, 1.25 mM NH_4_H_2_PO_4_, 1 mM KCl, 1x micronutrients (Murashige and Skoog Micronutrient Salts 100x, MSP18-10LT), in 0.8% agar with pH 5.7. The seeds were sterilized by soaking in 75% ethanol for 1 minute and 50% bleach for 5 minutes, 25% bleach with 0.2% Triton for 4 minutes, then rinsed with sterile water 8 times. After sterilization, seeds in the 2 mL Eppendorf tube were wrapped in tin foil and put into a 4°C fridge for cold stratification. Seeds were left in the fridge for 3 days for *A. thaliana* and *C. rubella*, or 5 days for *A. lyrata*. Cold stratification was not needed for efficient germination of *B. oleracea* seeds. We staggered the planting of different species to account for different germination times of the four plant species, allowing us to harvest seedling root and shoot tissues of plants at the same 5-7 day old growth stage of all species on the same day. *A. lyrata* was planted on the agar plates 3 days before *A. thaliana*, *C. rubella* was planted 1 day after *A. thaliana*, and *B. oleracea* seeds without cold stratification were planted 2 days after *A. thaliana*. The conditions of the growth chamber (Percival Scientific) were set to 16 hours light at 22°C and 140-150 umol/m^2^/s illumination followed by 8 hours dark at 19°C, with the light turning on at 5am. Root, shoot, and whole seedlings were sampled between 10am-12pm.

In order to enable sampling of mature leaf and flower bud samples on the same day of all four species, planting times were staggered and growth conditions were modulated to synchronize growth of all four species. For mature leaves, we sampled the youngest fully expanded rosette leaf from plants grown in soil in a plant grow room or cold room. *B. oleracea* mature leaves were harvested from plants that were two months old grown at 13°C under a light cycle of 12 hours light/12 hours dark. *A. lyrata* was three months old at harvest, grown at 4°C with a light cycle of 12 hours light/12 hours dark. *A. thaliana* and *C. rubella* were grown in conditions of 16 hours light at 28°C and 8 hours dark at 23°C and harvested at 2 months old. Mature leaves were sampled between 10am and 12pm.

For floral buds, we sampled young buds on the main stem from plants grown in soil, where the petal was just emerging (∼1 mm visible). All four plant species were grown at 16 hours light at 22°C and 8 hours dark at 19°C when floral buds were collected. *A. thaliana* and *B. oleracea* flowered at about two months old without cold treatment. To induce flowering in *A. lyrata* and *C. rubella*, two month old plants were transferred into a 4°C cold room for two months and then transferred back to conditions with 16 hours light at 22°C and 8 hours dark at 19°C for one month. In this way, all the four plant species were synchronized to flower on the same day and sampled between 10am-12pm.

*S. bicolor* Seeds were sterilized and planted vertically on agar plates as above, and placed in a growth chamber (Percival Scientific) set to 28°C with 800 µmol/m²/s illumination 16 hours light/8 hours dark, with the light turning on at 8am daily. Samples from 3 day old seedlings were collected between 10am-11am.

#### Nuclei isolation

Plant tissue samples were either processed immediately after harvest or flash frozen in liquid nitrogen and stored at −80°C for up to one year before nuclei isolation. Nuclei isolation from tissues was performed as follows.

Buffer 1 (lysis buffer) consisted of 0.275 M sorbitol (Sigma-Aldrich S6021), 0.1% Triton X-100 (Sigma-Aldrich 93443), 0.01 M MgCl_2_ (Ambion AM9530G), 1x protease inhibitor cocktail (Sigma-Aldrich 4693132001), 0.3 U/μl RNase inhibitor (Roche 03335399001), and 1 mM DTT (Teknova D9750).

Buffer 2 (wash and resuspension buffer) consisted of 1x phosphate buffer saline (without Mg and Ca), 1% BSA, 0.4 U/μl RNase inhibitor, and 1 mM DTT.

For isolation of nuclei from mature leaves containing high levels of chloroplasts, an extra modified wash and resuspension buffer (buffer 3) was used to eliminate chloroplasts before the final wash and resuspension step by buffer 2. It consisted of 0.275 M sorbitol, 0.2% Triton X-100, 0.2% NP40 (Thermo Fisher Scientific PI28324), 0.01 M MgCl_2_, 1x protease inhibitor cocktail, and 1 mM DTT.

All nuclei isolation steps were carried out in a cold room at 4°C. For each sample, 20-100 mg fresh or frozen tissue was chopped for 3 minutes using a razor blade on a glass plate with 100-200 μl buffer 1, then transferred to a petri dish with 1.5 mL cold buffer 1 and allowed to rest on ice with gentle shaking for 2 minutes. For multiplexed samples, tissues from different species were weighed separately before being combined and chopped together. The samples were then transferred to a 48-well filter plate (25 μm, 4 mL, Agilent, 201003-100), which was pre-wet with 1 mL cold buffer 1 before it was placed on the receiving plate attached to a QIAvac96 (Qiagen). Pressure was maintained below 100 bar during filtration. The filtered nuclei solution was centrifuged at 500x g for 10 minutes at 4°C to pellet the nuclei, after which the supernatant was discarded.

For samples with low or no chloroplast content: The pellet was resuspended by gentle flicking and pipetting, washed with 2 mL cold buffer 2, centrifuged at 500x g for 5 minutes, and the supernatant was discarded. For nuclei isolation from mature leaves with high chloroplast content: The pellet was resuspended by gentle flicking and pipetting and 10 mL cold buffer 3 was added, centrifuged at 500x g for 5 minutes, and the supernatant was discarded. The pellet was washed with 2 mL cold buffer 2, centrifuged at 500x g for 10 minutes, and the supernatant was discarded. For both protocols, pellets from the final centrifugation were resuspended in 100 μl cold buffer 2 for snRNA-seq, or cold 1x nuclei buffer (10X Genomics PN-2000207) for snATAC-seq.

Nuclei quality and quantity were assessed by flow cytometry and microscopy before continuing to the single nuclei partitioning and barcoding step. A BD Accuri C6 plus flow cytometer was used to analyze the quality and quantity of nuclei stained by propidium iodide (Sigma-Aldrich P4864), added to a final concentration of 50 μg/mL. The stained samples were incubated on ice in darkness for 5-10 min prior to analysis. The analysis was conducted using light-scatter and fluorescence signals produced by a 20 mW laser illumination at 488 nm. Signals corresponding to forward-angle scatter (FSC), 90° side scatter (SSC), and fluorescence were collected. Fluorescence signals (pulse area measurements) were screened using the following filter configurations: (a) FL-2, with a 585/40 nm band-pass filter, and (b) FL-3, with a 670 nm long-pass filter. Threshold levels were set empirically at 80,000 for FSC-H to exclude debris commonly found in plant homogenates. Templates for uni- (FL2-A and number of nuclei) and bi-parametric (FL2-A and FL3-A) frequency distributions were established. Upon identifying the region corresponding to nuclei, data was collected to a total count of 5,000–10,000 nuclei. The flow cytometer was operated at the Slow Flow Rate setting (14 μl sample/minute), with data acquisition for a single sample typically taking 3–5 minutes. Different plant species exhibited varying patterns of somatic endoreduplication within most of their tissues and organs. For example, in young *A. thaliana* roots and cotyledons, this variation is reflected in the form of multiple clusters within the distribution, corresponding to nuclei forming a 2C, 4C, 8C, 16C, etc., of the endoreduplicative series^61^. All the nuclei of all the endoreduplicative series were counted and divided by the actual volume analyzed in the flow cytometer to obtain the concentration of the nuclei. An EVOS M5000 microscope was also used for validation of nuclei quality and quantity by analyzing 7 μl diluted nuclei sample stained by 1 µl of diluted SYBR Green dye (Thermo Fisher Scientific S7585 diluted 1:10,000 in water) using a C-Chip Disposable Hemocytometer. For nuclei prepared from fresh tissue samples, most of the nuclei in each sample showed well-resolved edges without obvious evidence of blebbing. In the case of nuclei prepared from frozen samples, where cell debris around the nuclei can interfere with these microscopy observations, nuclei quality was primarily assessed by comparing to fresh nuclei’s distribution and the endoreduplicative series pattern on the flow cytometer.

#### Single nuclei library preparation

Single nuclei RNA-seq libraries from *A. thaliana*, *A. lyrata*, *C. rubella*, and *B. oleracea* tissues (two flower bud, two mature leaf, two seedling shoot, and 4-7 seedling root samples for each species, plus four whole seedling samples for *A. thaliana* and *B. oleracea*, see **Supplementary Table 5**), were prepared as follows. The nuclei suspension was diluted to a concentration of 700-1000 nuclei/μl using cold buffer 2. Single nuclei were partitioned and barcoded using a Chromium system (10x Genomics) with the Chromium Next GEM Single Cell 3’ Reagent Kits v3.1 (Dual Index). Approximately 20,000 nuclei in total were loaded for single species samples, and up to 60,000 nuclei for multiplexed species samples. Library creation was carried out following the manufacturer’s protocol. Sequencing was performed on an Illumina NovaSeq 6000 S4 flow cell or NovaSeq X plus 10B or 25B flow cell, using 2×150 bp paired-end sequencing, targeting 200M fragments per species for RNA-seq samples.

Single nuclei RNA-seq libraries from *S. bicolor* tissues (four *S. bicolor* root samples, three *S. bicolor* leaf samples, see **Supplementary Table 5**) were prepared as follows. Note that some samples were multiplexed on the same lane of the cartridge with samples from several other plant species, which were resolved independently by mapping to their respective reference genomes after sequencing. For each sample, the pooled nuclei suspension was diluted to a concentration of 600-900 total nuclei/μl using cold buffer 2. Single nuclei were partitioned and barcoded on a BD Rhapsody system (BD Rhapsody HT Xpress, BD Rhapsody Scanner), using the BD Rhapsody WTA V3 kit and cartridge, loading 50,000-150,000 total nuclei per lane. Library creation was carried out following the manufacturer’s protocol. Sequencing was performed on an Illumina NovaSeq X plus 25B flow cell, using 2×150 bp paired-end sequencing and targeting an average of at least 50,000 reads (25,000 fragments) per nucleus.

### DAP-seq Analysis

#### Primary DAP-seq data analysis pipeline

Fastq files were quality filtered and adapters were trimmed using BBTools v38.96 bbduk.sh^62^ with parameters k=21 mink=11 ktrim=r tbo tpe qtrim=r trimq=6 maq=10, and then aligned to the corresponding reference genome using bowtie2 v2.4.2^63^ and parameters --no-mixed --no-discordant. Bam files from negative control wells were merged using the samtools v1.15.1^64^ merge command, to generate a single background file for each 96-well plate. The MACS3 v3.0.0a6^65^ callpeak command was used to generate narrowPeak files, using the merged background as the control, with the parameters --call-summits --keep-dup 1 and –gsize with the total fasta genome size. Motifs were called using MEME suite v5.3.0^66^ command meme with parameters -dna -revcomp -mod anr -nmotifs 2 -minw 8 -maxw 32, and with a background model (-bfile) generated from the entire reference genome file with fasta-get-markov -m 0 -dna.

The resulting datasets were filtered to exclude those TFs and replicates with poor performance in the DAP-seq assay. As a measure of successful binding site enrichment, we calculated the fraction of reads in peaks (FRIP) scores and only used data from DAP-seq assays that generated FRIP scores of at least 0.05. For experiments including multiplexed species, we only considered those assays where all of the multiplexed species independently produced FRIP scores of 0.05 or greater. We found that multiplexing larger sets of species with varying genome sizes in a single multiDAP experiment sometimes resulted in large differences in read coverage between different TFs and in some cases differential enrichment of species. To mitigate the effect of low coverage resulting in false negatives in the peak-calling step (and thus lowering apparent c scores), we only used datasets that yielded at least 200,000 read pairs (400,000 reads) for each of the multiplexed species.

#### Assignment of DAP-seq peaks to target genes and conservation scores (c scores)

DAP-seq peaks files generated by the above analysis pipeline were further processed to identify target genes for each TF and in each species using a set of custom scripts. For each species, the corresponding gene annotation file (gff format) was filtered to retain only features corresponding to a single transcript per gene, for which the primary (or longest) transcript was selected. The script 1_assign_peaks_to_gff_features.sh (a wrapper for bedtools^67^ v2.31.0) was used to assign peaks to mRNA, exon, and CDS features individually. The output was further processed using the script 2_annotate_peak_targets.py, which integrates information from the different assigned feature types to assign each peak to up to two gene targets and categorize their location with respect to the target gene as either upstream, downstream, within the 3’ or 5’ UTR, within the CDS, or intronic. Peaks falling in intergenic regions between two divergent genes were assigned to both flanking gene targets, while peaks falling within the transcript (UTR 5’, CDS, intron, or UTR 3’) were assigned to only a single gene target.

The output of the above scripts was further processed with the script 3_filter_annotated_peaks_and_calculate_c_scores.ipynb, which was used to filter for peaks falling within the regions spanning −2000 bp to +500bp with respect to the target gene’s start codon. Peaks were only considered if they had a strength of at least 5-fold over background (as defined in the 7th signalValue column of the narrowPeak file format), and if they fell in one of the following regions: upstream, 5’ UTR, CDS, or intron. Each target gene was then mapped to an orthogroup, as defined by applying OrthoFinder v2.5.5^5^ with default parameters to the protein sequences of all species. To improve orthogroup resolution within the more closely related species, separate tables of orthogroups were generated and used to analyze the four species datasets and the 10 species datasets. Finally, a conservation score (c score) was assigned to each TF-orthogroup assignment, representing a count of species in which the given TF also targets at least one gene within the same orthogroup. To exclude non-nuclear genes from the analyses, orthogroups containing one or more *A. thaliana* genes originating from the chloroplast or mitochondrial chromosomes were ignored.

For comparative analyses of the brassica and grass lineages, we used OrthoFinder with a manually curated species tree to call hierarchical orthogroups at each node in the species tree. To identify lineage-specific TF target genes within the brassica and grass families, we used the hierarchical orthogroups called at the base node of the corresponding clade.

See code availability statement for scripts, reference genome files, and sample data files.

#### DAP-seq signal comparisons

For assessment of DAP-seq signal similarity across individual DAP-seq assays, we compared bigwig files using deeptools^68^ v3.5.2 multiBigwigSummary bins command with parameter “--binSize 50”. In order to reduce computation time, this analysis was restricted to only the first chromosome of the *A. thaliana* genome using the flag “--region Chr1”. The resulting output was processed to generate a correlation matrix with the deeptools command plotCorrelation and “--corMethod pearson”.

#### GO term enrichment

We used clusterprofiler v4.6.2^69^ to perform gene ontology (GO) enrichment analysis on the list of target genes for each TF and c score. We first converted the TAIR gene names into the Entrez ids. Then we calculated enrichment values for Biological Processes and filtered for enriched terms with level six to avoid broad GO terms. We then used the “simplify” function to combine terms with a similarity cutoff of 70%.

#### Nucleotide diversity in TFBSs

We used the 1001 genomes polymorphism data^26^ to calculate nucleotide diversity (pi) with vcftools v0.1.17^70^. We only considered regions flanking peak summits (+/− 30 bp from the peak summit location) that were located within 2000 bp upstream of the assigned gene’s coding sequence start. We did not include regions downstream of the start codon or regions overlapping coding sequences in the 2000 bp upstream of the assigned gene.

### snRNA-seq analysis

#### snRNA-seq atlas construction

Raw fastq files for each library were filtered using BBTools v38.96 bbduk.sh^62^ to remove read pairs with 31-mers in read 2 matching common ribosomal 31-mers. Nuclear reference genomes for all species were downloaded from Phytozome where available, and supplemented with organellar genomes from NCBI. For brassica species (profiled with 10X Chromium), rRNA-filtered reads were pre-processed with Cellranger count v7.0.1 and the resulting bam file was run through Velocyto^71^ to generate spliced and unspliced count matrices. Sorghum libraries (profiled with BD Rhapsody) were pre-processed with the BD Rhapsody CWL pipeline (v2.2, available at bitbucket.org/CRSwDev/cwl) and in parallel pre-processed with STAR with parameter ‘*--soloFeatures Velocyto’* to generate spliced and unspliced count matrices. After pre-processing, QC metrics were compiled for each sample by (1) calculating the total UMIs and genes from each cell barcode (CBC), (2) calculating the proportion of UMIs from each CBC mapping to organellar genomes, (3) associating each CBC with an overall rate of unspliced vs. spliced transcripts, (4) running the R package diem^72^ to estimate the rate of ambient RNA in each CBC. Cell barcodes were then filtered to those with at least 400 nuclear UMIs, 200 nuclear genes, 90% nuclear UMIs, 10% unspliced UMIs, and debris score <=2. The 400 UMI threshold was relaxed to 300 UMIs for flower bud samples in order to obtain enough cells for meaningful clustering, but all other filters remained unchanged. In some experiments nuclei from multiple species were pooled together, as inter-species sequence divergence is sufficient to distinguish transcripts based on genome alignment, and we did not observe any reads multi-mapping to annotated genes from different species. For these mixed-species samples, additional filtering was performed by defining background proportions for each species using all UMIs in the raw expression matrix, and then running a chi-squared test for each CBC to determine whether the species proportions calculated from the CBC’s associated UMIs were significantly different (p<0.01) from background. CBCs passing this test were assumed to contain nuclei from one or more species. To determine which species were present, a per-species UMI threshold was defined as the 99th percentile of per-species UMIs across all CBCs failing the chi-squared test. Each CBC was then assigned an estimated ambient RNA proportion by comparing total UMIs from ‘present’ species to total UMIs from all species, and CBCs with >50% estimated ambient RNA were further filtered from downstream analysis. As the rate of unspliced transcripts proved to be an especially informative metric^73^ for sorghum libraries, these libraries were further filtered by detecting the inflection point (aka ‘knee’ in a knee-cliff plot) for all CBCs based on unspliced rate, and only keeping barcodes with a rate exceeding this threshold.

After filtering each sample, count matrices for all remaining CBCs with organelle genes removed were loaded into Seurat (v5.0.0)^74^, and *SCTransform* v2 normalization was performed, followed by an initial clustering using Seurat’s *RunPCA* (with 5,000 variable genes and 20 PCs), *FindNeighbors*, and *FindClusters* (resolution 0.8). Next, marker genes were identified for each cluster using *FindAllMarkers* (with *logfc.threshold = .5, min.pct = .4, only.pos = T)* filtered to those with *p_val_adj<0.05*, and clusters with zero marker genes and significantly lower UMI counts than other clusters were removed as low quality clusters. Next, DoubletFinder^75^ was run on remaining CBCs and those with doublet score >0.4 were removed. To prepare final atlases, samples from the same species and tissue (seedling, leaf, or flower bud) were merged into a single seurat object, which was then split by sequencing batch and re-normalized with SCT per batch before performing a final PCA (50 PCs) and clustering (resolution=1.0 for brassica and sorghum leaf, resolution=2.0 for sorghum root).

For *A. thaliana* libraries, we assigned cells to unique cell types and developmental stages via label transfer from three well-annotated large-scale atlases (a root atlas^28^, leaf atlas^29^, and a 6-day old seedling atlas^30^) as well as cluster-specific enrichments of known marker genes^31^. Labels were transferred from each published reference atlas to each SCT-normalized batch of the *A. thaliana* seedling and leaf datasets with Seurat’s *FindTransferAnchors* function using 20 PCs, and each CBC was putatively annotated with the highest scoring label. Additionally, marker genes from each cluster in each of the three *A. thaliana* tissue datasets were tested for enrichment (hypergeometric enrichment test) of a large suite of previously computed marker genes downloaded from scPlantDB^31^. Final cell type labels were compiled from these two sources, with additional manual annotation based on literature support for data-derived markers when there were discrepancies or no significant matches.

As a means of validating clustering and cell type labeling, we directly integrated our 35,169 *A. thaliana* seedling snRNA-seq transcriptomes with an external *A. thaliana* snRNA-seq dataset^32^ derived from 7-day old seedling roots, resulting in a combined dataset of >45,000 nuclei. To combine in-house and external samples, and samples from different tissue subtypes (root, shoot, and whole seedling), we tested a number of different integration techniques (CCA, RPCA, and Harmony) using Seurat’s *IntegrateLayers* function, and evaluated each as to the separation of root- and shoot-derived CBCs and mixing of internal and external root-derived CBCs. The RPCA method using a sample tree to fix the integration order proved to be the most appropriate, with root-specific cell type labels overwhelmingly composed of nuclei derived from root and whole seedling samples (for example, trichoblast-labeled cluster 29 log2 root:shoot ratio: 5.08, see **Extended Data Fig. 3a**), and comprising an approximately even mix of nuclei from the internal and external root datasets (cluster 29 log2 internal:external ratio: −0.65). Conversely, clusters receiving leaf-specific labels were strongly depleted for root-derived nuclei (including all nuclei from the external root dataset) (for example mesophyll-labeled cluster 13, with log2 root:shoot ratio: −5.24). Integration also revealed mixed internal-external clusters (including clusters 35, 17, and 19) that appeared to be associated with endodermis subtypes not typically captured by protoplast-based assays. These clusters shared expression of top endodermis-specific genes (for example AT3G32980/PRX32; see **Extended Data Fig. 3c**) but had distinct expression profiles. Cells in cluster 35 were mostly labeled mature endodermis via label transfer from the protoplast-based reference root atlas (see **Extended Data Fig. 3a**), yet unlike other endodermis clusters expressed key genes associated with lignin biosynthesis, including CASP1 and ERK1. Cells in cluster 17 received mixed low-confidence labels from the root atlas label transfer, and strongly expressed GELP96 and CYP86A1, genes with known roles in suberin biosynthesis and polymerization. The vast majority of external root nuclei in this cluster were labeled ‘suberized-endodermis’ in the original publication. The publication noted that this cell type was only detected in single nuclei rather than single cell datasets, potentially due to difficulty in digesting suberized cell walls. Cells in cluster 19 received low confidence atlas labels as well, yet included external cells from unlabeled ‘cluster 14b’ in the original publication. This cluster was also noted in the publication as appearing only in snRNA datasets, and showed tentative evidence of a role in cortex development and root epidermal patterning via expression of SCRAMBLED/STRUBBELIG. Both internal and external cells in our cluster 19 were differentiated from other endodermis cells by strong expression of FACT, a HXXXD-type acyl-transferase, and TLL1, a triacylglycerol lipase, both involved in synthesis of root wax components. A recent study^76^ identified TLL1 as a strong marker of suberized phellem in *A. thaliana,* and thus we labelled this unique cluster ‘phellem’. Clustering of internal *A. thaliana* seedling cells without integration of the external dataset yielded a cluster with very similar expression. We note that a cluster was also identified in a separate published nuclei-based dataset^77^ that showed highly similar expression patterns and was described as an ‘endodermis cluster with epidermis precursor cells’. Additionally, we found a distinct subset of cells in each of our three other brassica seedling atlases with a best match to cells in cluster 19, indicating that this cell type is not unique to *A. thaliana*.

Aside from the *A. thaliana* seedling atlas, all other atlases were constructed by merging relevant libraries, as no batch effects were apparent and integration was not necessary. Final cell type labels from the *A. thaliana* seedling, leaf, and flower atlases were transferred to the respective datasets from the other brassica species using SAMap^33^, selecting the top-scoring *A. thaliana* label per cell. Cell type labels for sorghum atlases were assigned based on the expression of key experimentally-validated cell type marker genes found in published literature (see **Supplementary Table 9** and **Extended Data Fig. 7**), as well as enrichments for maize marker gene sets identified in previously published single-cell atlases^31^. After final cell type labels were assigned in each atlas, the top 100 marker genes per cell type were identified by filtering the results of Seurat function *FindAllMarkers* (with *assay=SCT*, *min.pct = 0.1, only.pos = T)* to those with *p_val_adj<0.01* and sorting by avg_log2-fold change.

#### TF-target co-expression analysis

SCT-normalized count data from the *A. thaliana* seedling atlas was used to calculate Spearman correlation coefficients between the expression profiles of each assayed TF and every other gene across all nuclei. To compute final TF-target co-expression scores, we compared the squared correlation coefficient for a true TF-target pair to the squared correlation coefficient for the TF and a matched control gene. To find an appropriate control gene, all genes in the genome were assigned to one of 10 equally sized bins based on their average expression across all nuclei (*expression-bin*), as well as a bin representing the number of species (out of 4) in which an ortholog was present (*ortho-bin*). We then compiled a list of 500 randomly chosen expressed target genes per TF, and identified a matched non-target gene for each from the same *ortho-bin* and *expression-bin.* Next each TF’s target genes were additionally binned by fold-change of the strongest DAP peak (5 equally sized *peak strength bins* per TF) and c score, and we calculated the average fold-change between the squared correlation coefficient of true target genes vs. matched non-target controls within each group. We repeated this procedure 100 times to obtain mean and standard error values for each group.

#### Cell type specificity and enrichment analyses

In order to calculate target gene summary scores, atlases for untreated seedling, leaf, and flower bud tissues of each species were first merged together. Target gene expression summary scores were computed for each TF’s expressed c1 through c4 target gene sets in each cell of each atlas using Seurat’s *AddModuleScore* function with *assay=”SCT”* and default parameters. Average summary scores for each cell type were visualized as a heatmap overlaid onto diagrams of plant tissue physiology using ggPlantmap^78^. Summary scores for seedling cells were also used to calculate an overall measure of ‘cell-type specificity’ for each TF’s target sets, by extracting the F statistic from an anova test using R function *aov* with formula *score ∼ celltype*. F statistic distributions across all TFs were plotted as boxplots for each c score and each species, and paired one-sided t-tests (using R function *t.test*) were performed for each species comparing F statistics from c4 vs. c3 TF target sets.

To compute enrichment or depletion of the expression of a TF’s c4 target genes in each cell type, separate one-sided Wilcoxon Rank Sum tests were run to test for significantly higher or significantly lower target expression summary scores for cells with a given cell type label compared to all other cells, and magnitude of the difference was calculated using the function *cohen.d* from the R package *effsize*. The same process was used to calculate TF target gene enrichment scores (aka TF activity scores) in each sorghum and *A. thaliana* cell type using subsets of brassica-c4 or grass-c2 target genes from the 10 species multiDAP dataset.

## Extended data figures and tables

**Extended Data Fig. 1.**
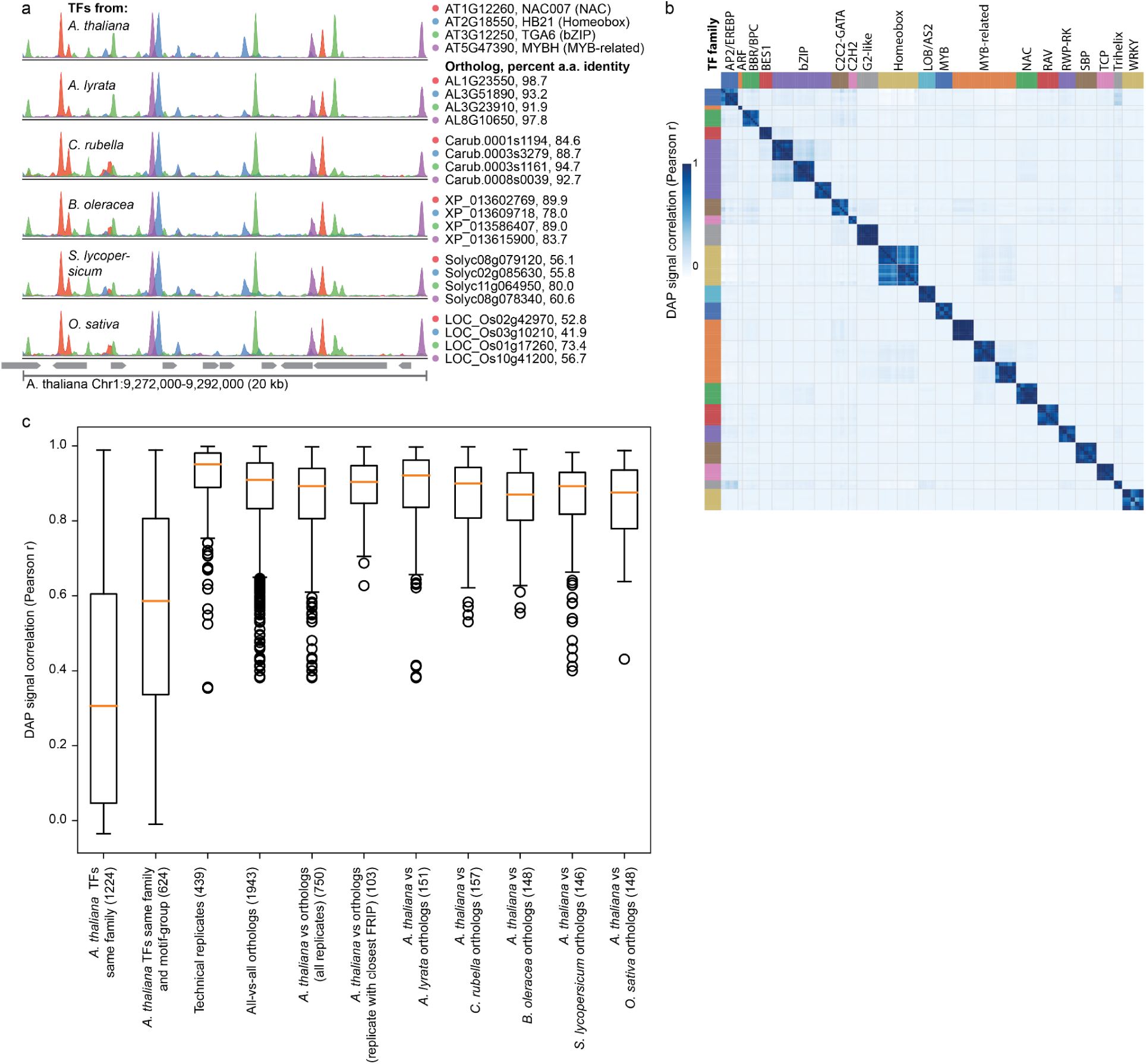
(a) Example 20 kilobase region of the *A. thaliana* genome shows similar TF-binding profiles of four example *A. thaliana* TFs and their corresponding orthologs from five other species. Numbers after TF IDs indicate percent amino acid identity compared to the *A. thaliana* ortholog. (b) All-vs-all genome-wide correlations of DAP-seq binding profiles between TFs from six species. Gray grid lines delineate individual groups of orthologous TFs and colored bars indicate TF families. (c) Boxplots showing median +/− interquartile range of correlations between different subsets of DAP-seq binding profiles. Numbers in parenthesis indicate the number of pairwise comparisons in each category. FRIP = fraction of reads in peaks, which serves as a measure of signal-to-noise in each DAP-seq library and can be used as an estimate of overall performance of the assay. Performance in the assay is variable between different TFs, and also varies slightly between technical replicates of the same TF. The category labeled “replicate with closest FRIP” partially controls for this confounding effect on correlations: for each comparison between an *A. thaliana* TF and an orthologous TF, one technical replicate of each TF was selected such that the difference in FRIP scores between the pair being compared was minimized. In the category labeled “same motif group”, motif-groups refers to groups of *A. thaliana* TFs that fall within the same TF family and also show highly similar motifs (see Table S1).

**Extended Data Fig. 2.**
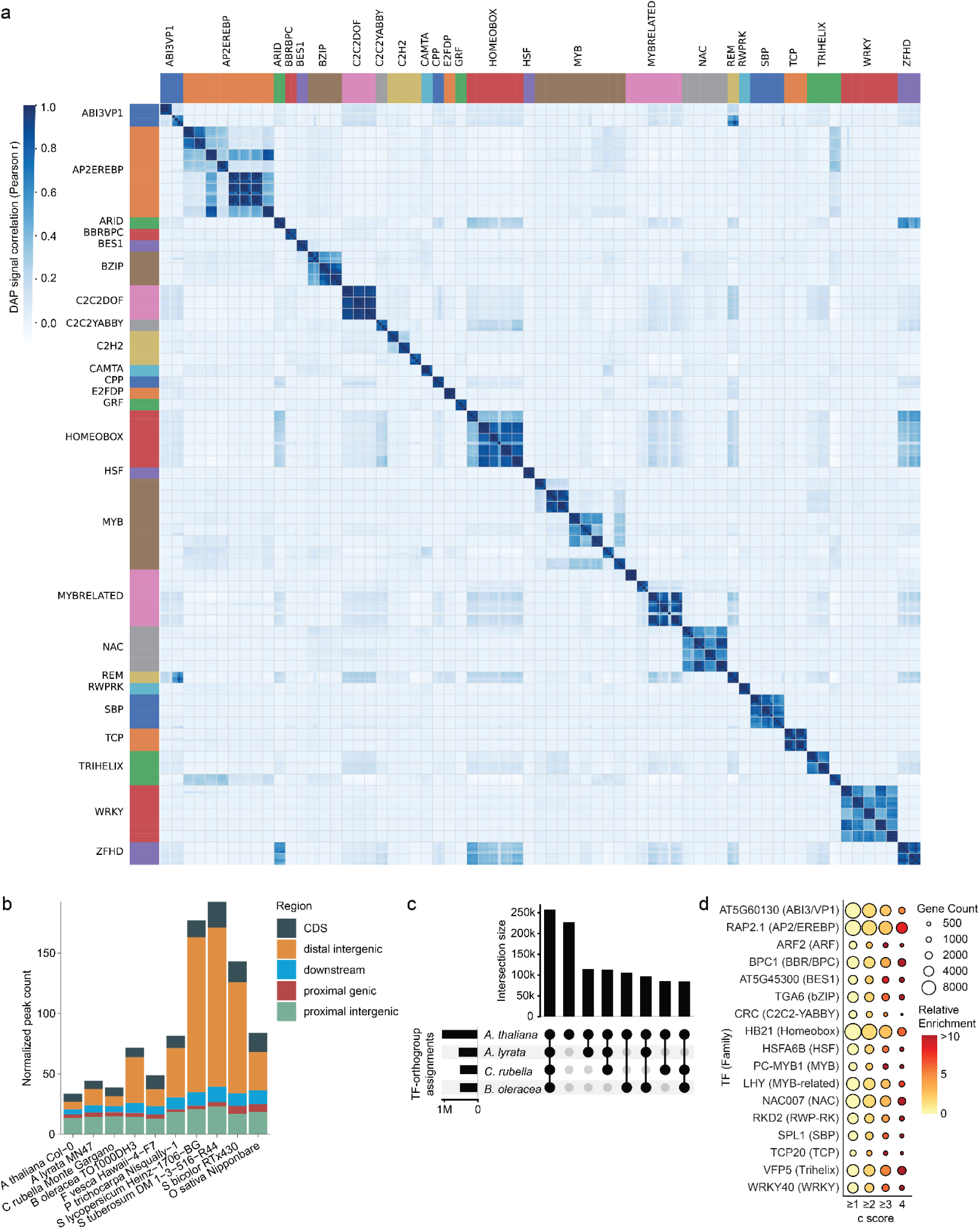
(a) All-vs-all genome-wide correlations of DAP-seq binding profiles between four replicates for each TF using either 1, 4, or 10 multiplexed species. Gray grid lines delineate individual TFs and colored bars indicate TF families. Mean correlation between single species replicates = 0.94, standard deviation = 0.08, n = 116; mean correlation between single and four species experiments = 0.94, standard deviation = 0.06, n = 211; two-sample t-test p = 0.56. Mean correlation of 10 species vs. single species experiments=0.87, standard deviation = 0.1, n = 163. (b) Count of peaks in genomic regions normalized by the number of total genes per species. CDS: coding sequence; distal intergenic: >2000 bp upstream of any start codon; downstream: within 2000 bp downstream of a stop codon; proximal genic: within 500 bp downstream of a start codon; proximal intergenic: within 2000 bp upstream of a start codon (c) Number of TF target genes shared between *A. thaliana* and other brassica species. (d) Overrepresentation test for conserved sites. A subset of representative example TFs are displayed. All 244 TFs were significantly enriched (p<0.01) at c scores ≥ 2.

**Extended Data Fig. 3.**
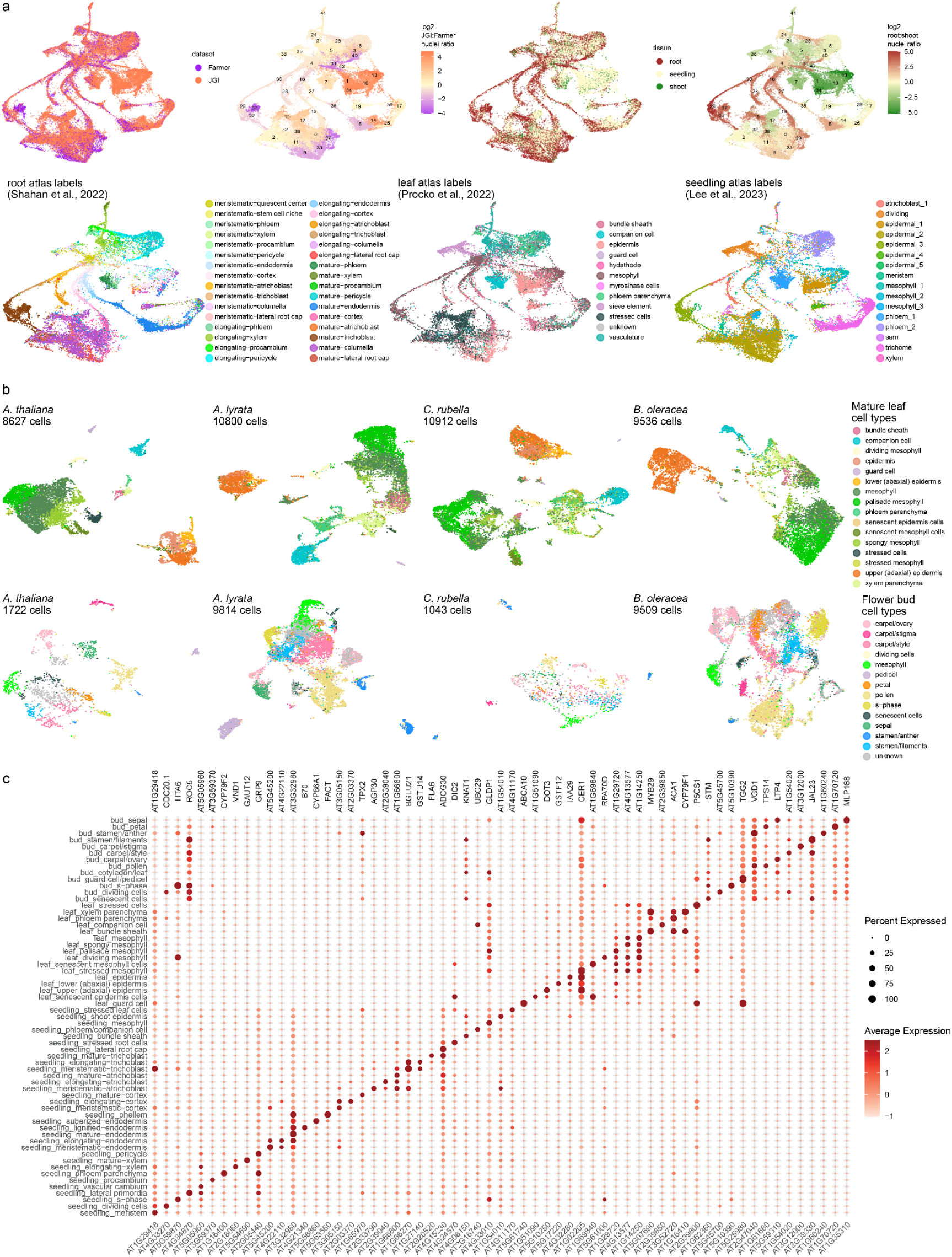
(a) Integration of internal and external *A. thaliana* seedling snRNA-seq datasets (first row) and results of label transfer from three reference atlases (second row). Only cells receiving a label with confidence score >0.4 are shown in these plots. (b) UMAP representations of mature leaf and flower bud snRNA-seq atlases from four profiled brassica species. (c) Dotplot of the top marker gene for each cell type, where dot size represents the percent of all nuclei with a specific cell type label showing non-zero expression of the gene, and color represents average normalized and scaled expression across these nuclei.

**Extended Data Fig. 4.**
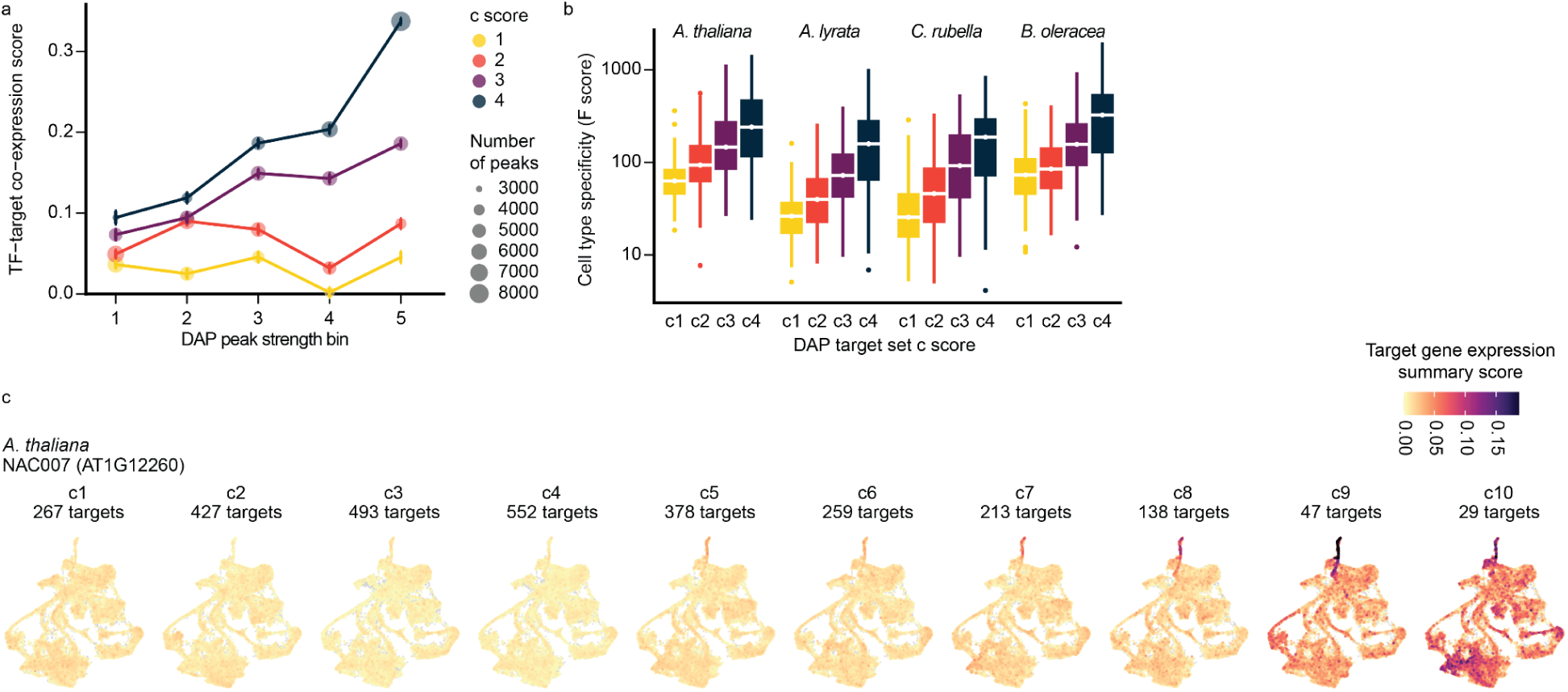
(a) TF-target co-expression across all nuclei in the *A. thaliana* seedling atlas as a function of multiDAP TF binding affinity and multiDAP peak conservation score. Y-axis represents the mean fold-change between true correlation of normalized TF expression with target expression vs. correlation with a matched control target gene. Point size represents the number of DAP peaks considered, and error bars represent standard error of the mean. (b) Boxplots showing median +/− interquartile range of cell type specificity scores as a function of target conservation for 244 TFs in each of the four brassica seedling snRNA-seq atlases. Cell type specificity was defined as the F-statistic from an ANOVA test of the extent to which per-nucleus target gene expression varied systematically across labeled cell types. All distributions within a species were significantly different from each other (FDR < 0.05) using paired one-sided t-tests. (c) NAC007/VND4 target gene expression summary scores using target c scores from extended 10-species multiDAP dataset. Note that two target genes are excluded as they were not expressed in any cells in the snRNA-seq dataset.

**Extended Data Fig. 5.**
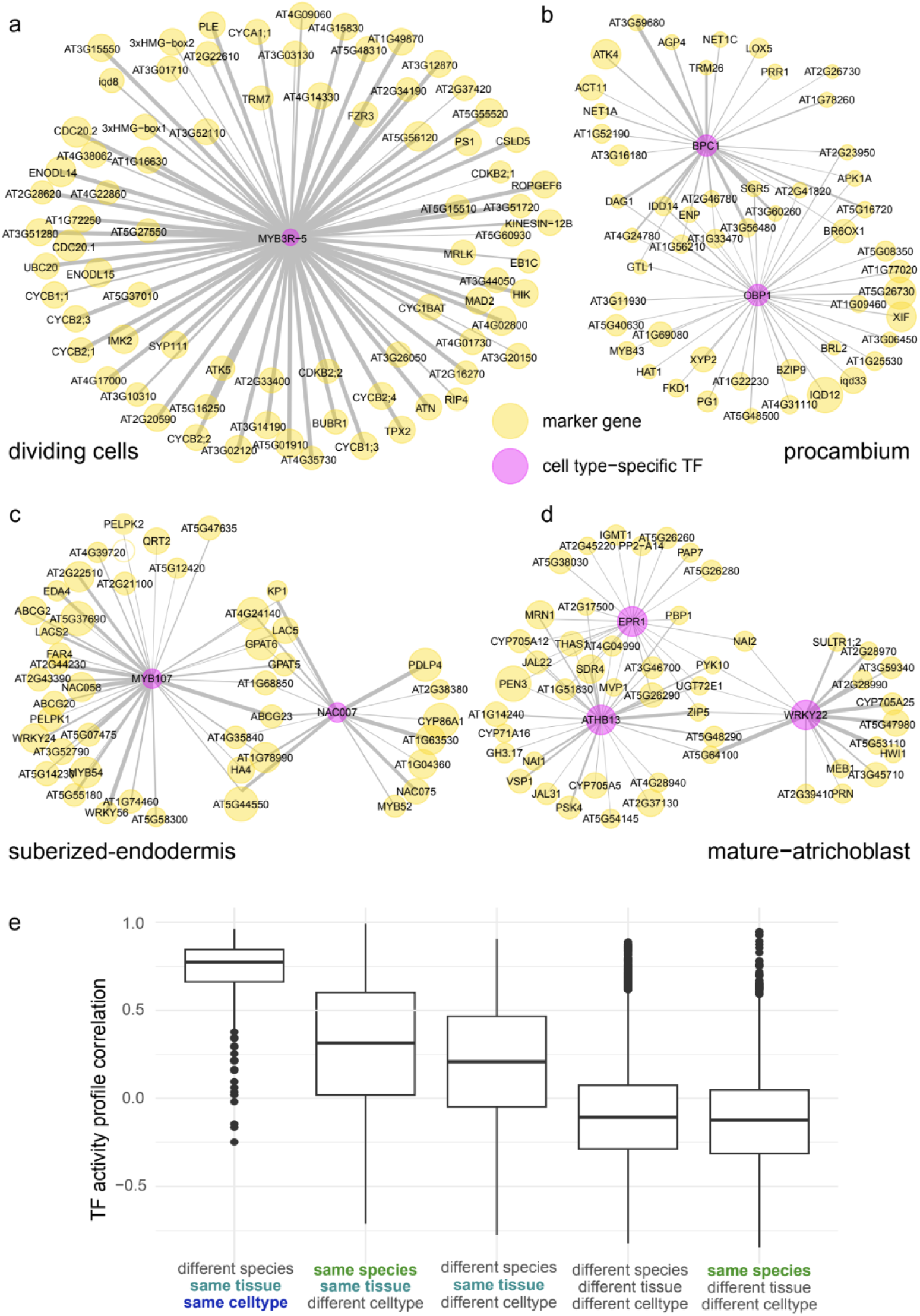
Network diagrams showing TFs with enriched activity scores (based on summarized expression of all c4 target genes) and the cell type-specific marker genes they target for four example cell types: a) dividing cells, b) procambium, c) suberized-endodermis, and d) mature-atrichoblast. TFs are shown in pink and marker genes are yellow, with a gray line connecting them if a brassica-c4 TFBSs for the TF was associated with the marker gene. Line thickness corresponds to relative binding strength (measured as DAP-seq peak height) and marker gene node size represents relative cell type specificity (measured as expression log2-fold enrichment).

**Extended Data Fig. 6.**
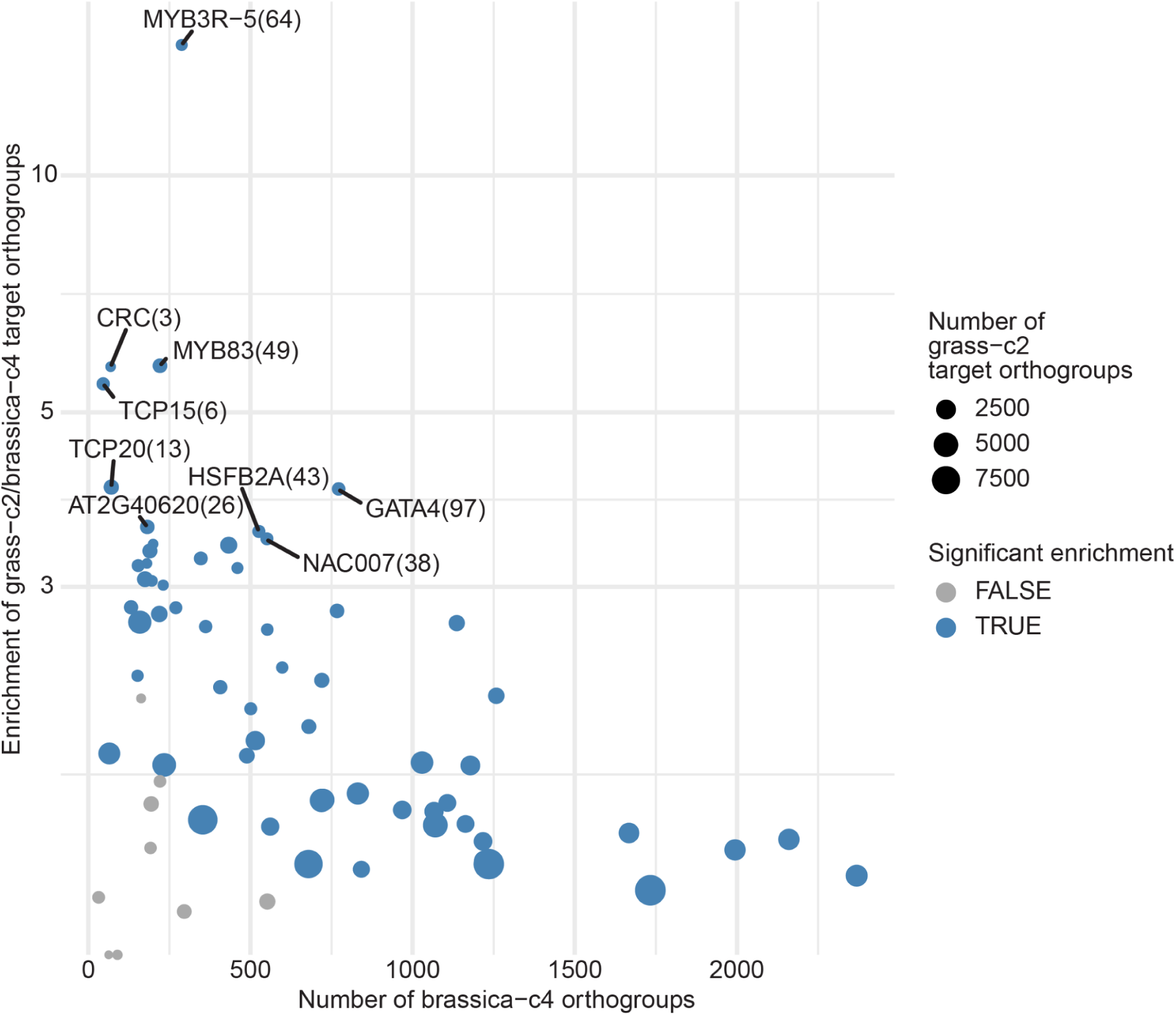
Scatterplot showing enrichment of each TF’s core regulon (i.e. target orthogroups conserved in both grasses and all four brassica), for TFs assayed in the 10 species multiDAP experiment. To ensure meaningful statistics, only the 70 TFs targeting at least 150 total brassica-c4 or grass-c2 target orthogroups were tested. Each point represents a single TF, and x-axis shows the total number of brassica-c4 orthogroups, dot size shows the total number of grass c2-orthogroups, and the y-axis shows the enrichment of orthogroups shared between these two sets. To calculate enrichments, true counts of core regulon target orthogroups for each TF (shown in parentheses after TF name for top examples) were compared to background distributions generated by shuffling orthogroup labels for c2-grass TFBSs 1,000 times. Significance (dot color) was calculated as the number of times the shuffled core regulon count met or exceeded the actual count, followed by FDR correction. Significance was assessed at FDR<0.05. Enrichment score (y-axis) was calculated as the fold change between the actual count and the average shuffled count.

**Extended Data Fig. 7.**
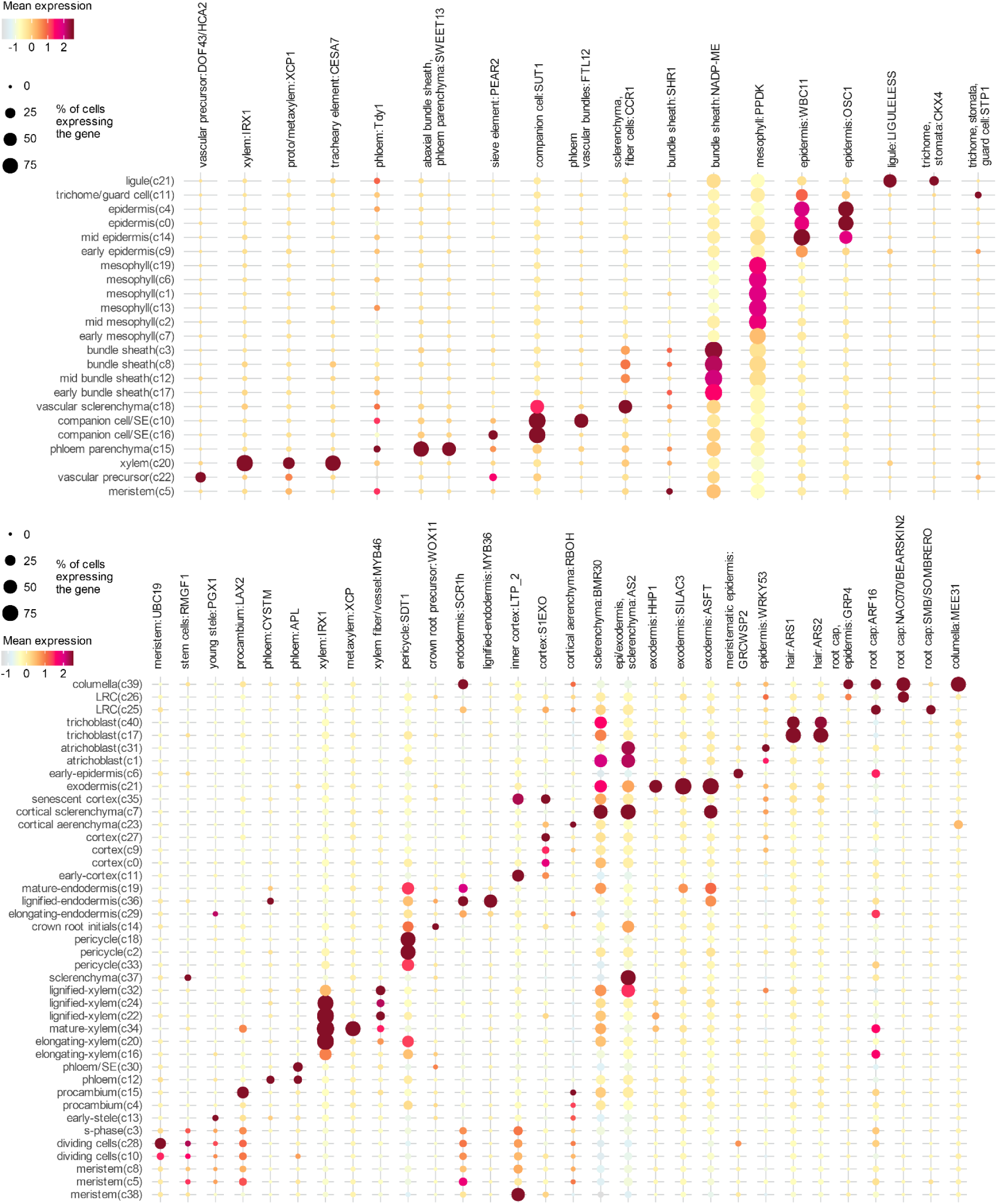
Dotplots showing expression of sorghum orthologs of experimentally validated cell type marker genes culled from published literature in each cluster of the sorghum leaf (top) and root (bottom) snRNA-seq atlas. Column labels specify the cell type(s) where gene expression was experimentally observed/validated, followed by the gene alias. LRC = lateral root cap. For gene IDs and publication references, see **Supplementary table 8**.

**Extended Data Fig. 8.**
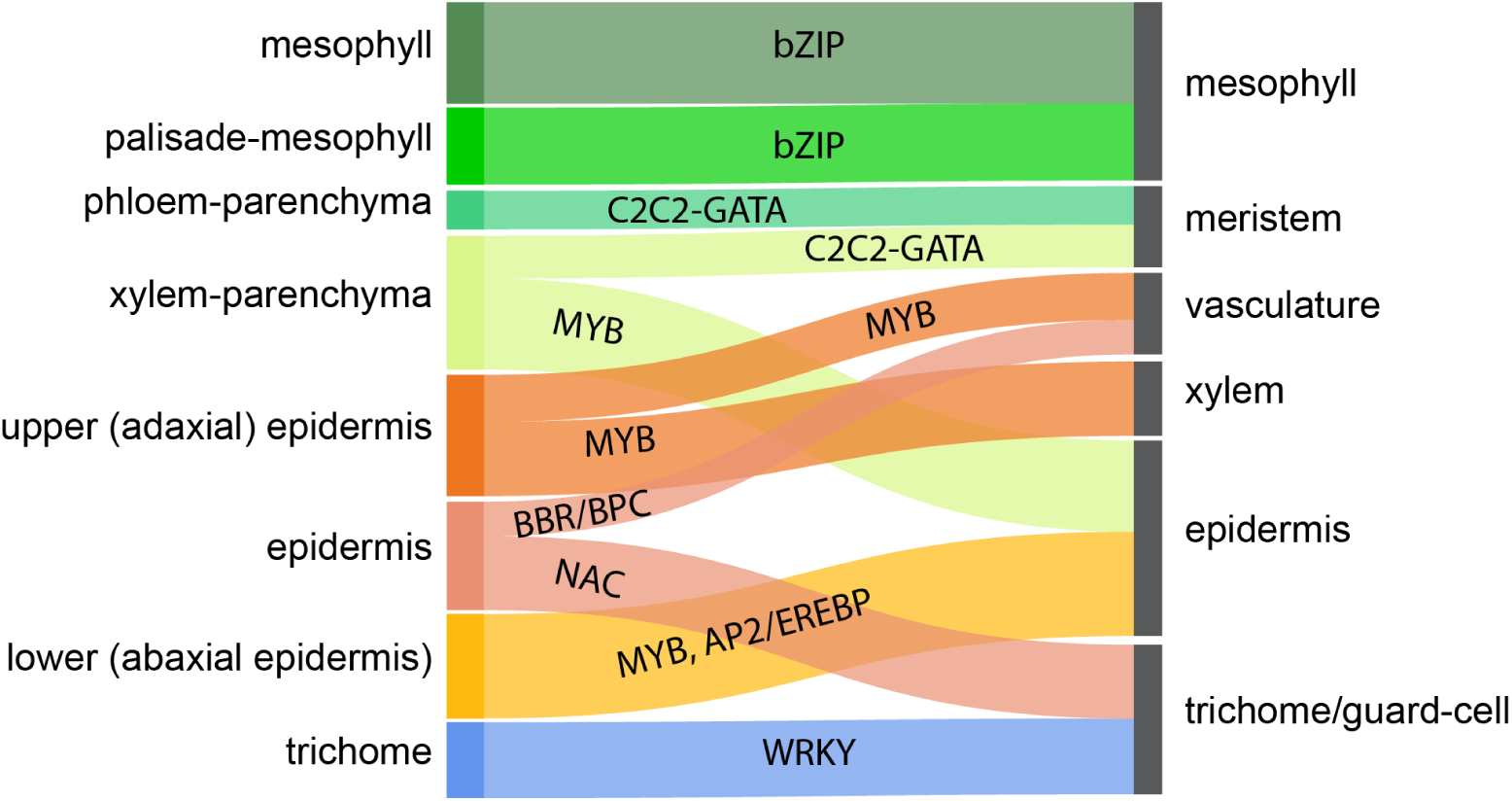
Sankey plot showing top activity score correlations for each sorghum leaf cell type (right) among all *A. thaliana* leaf cell types (left). A vector of TF activity scores was constructed for each sorghum and *A. thaliana* cell type, quantifying enrichment of each TF’s core regulon target genes (i.e. conserved in both grasses and all four brassica) using Cohen’s d. Then Pearson correlations were calculated between the TF activity score vectors of all cross-species cell type pairs. Ribbons in the diagram represent the top two strongest correlations for each sorghum cell type, after filtering to correlations with r>0.3. Ribbon width corresponds to Pearson r^2^ value, and ribbons are labeled with TF families found in the top 5 enrichments for both the *A. thaliana* and sorghum cell type.

**Extended Data Fig. 9.**
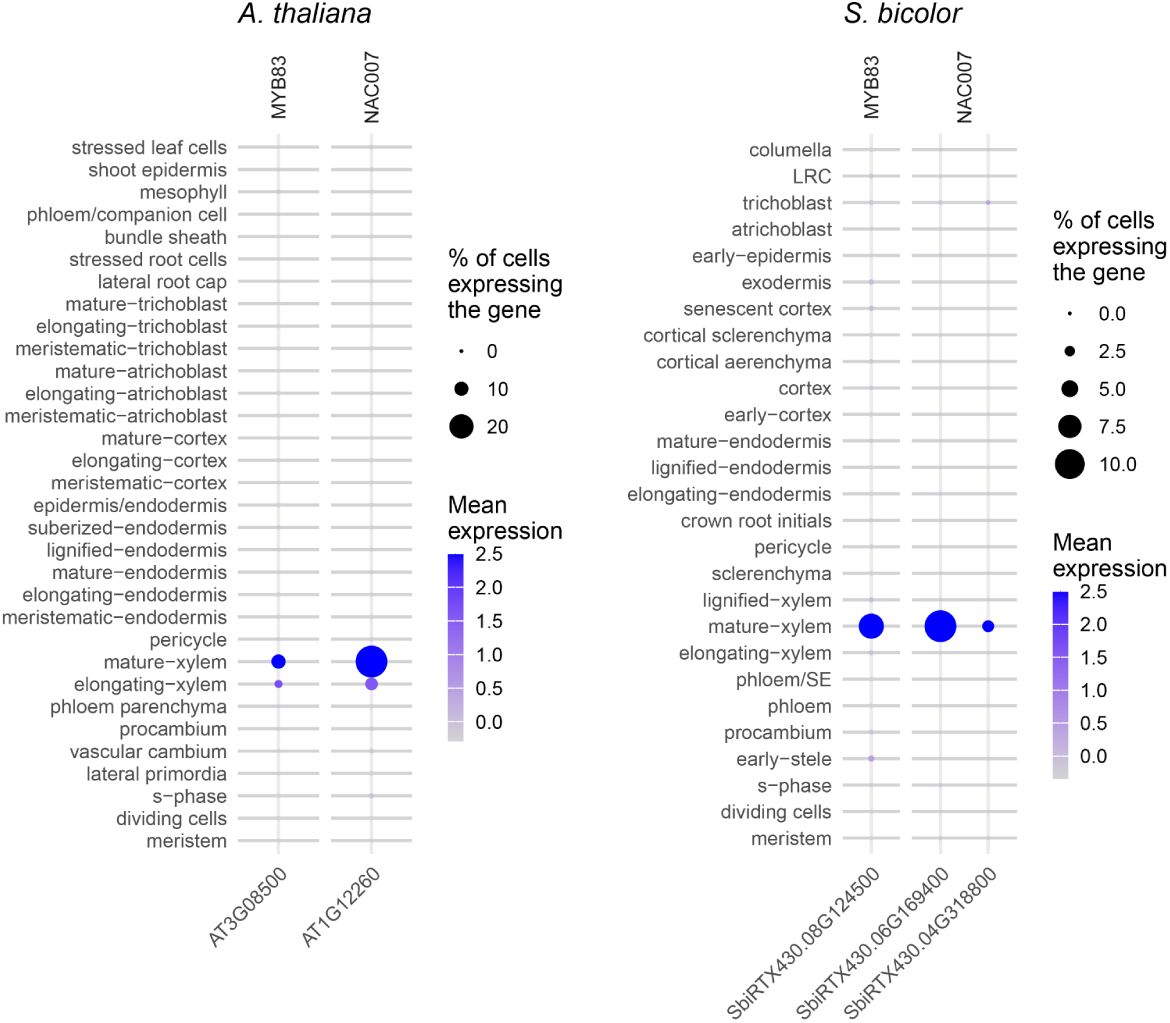
Dotplots showing expression of TFs MYB83 and NAC007 across all *A. thaliana* seedling cell types and all sorghum root cell types.

**Extended Data Table 1.**
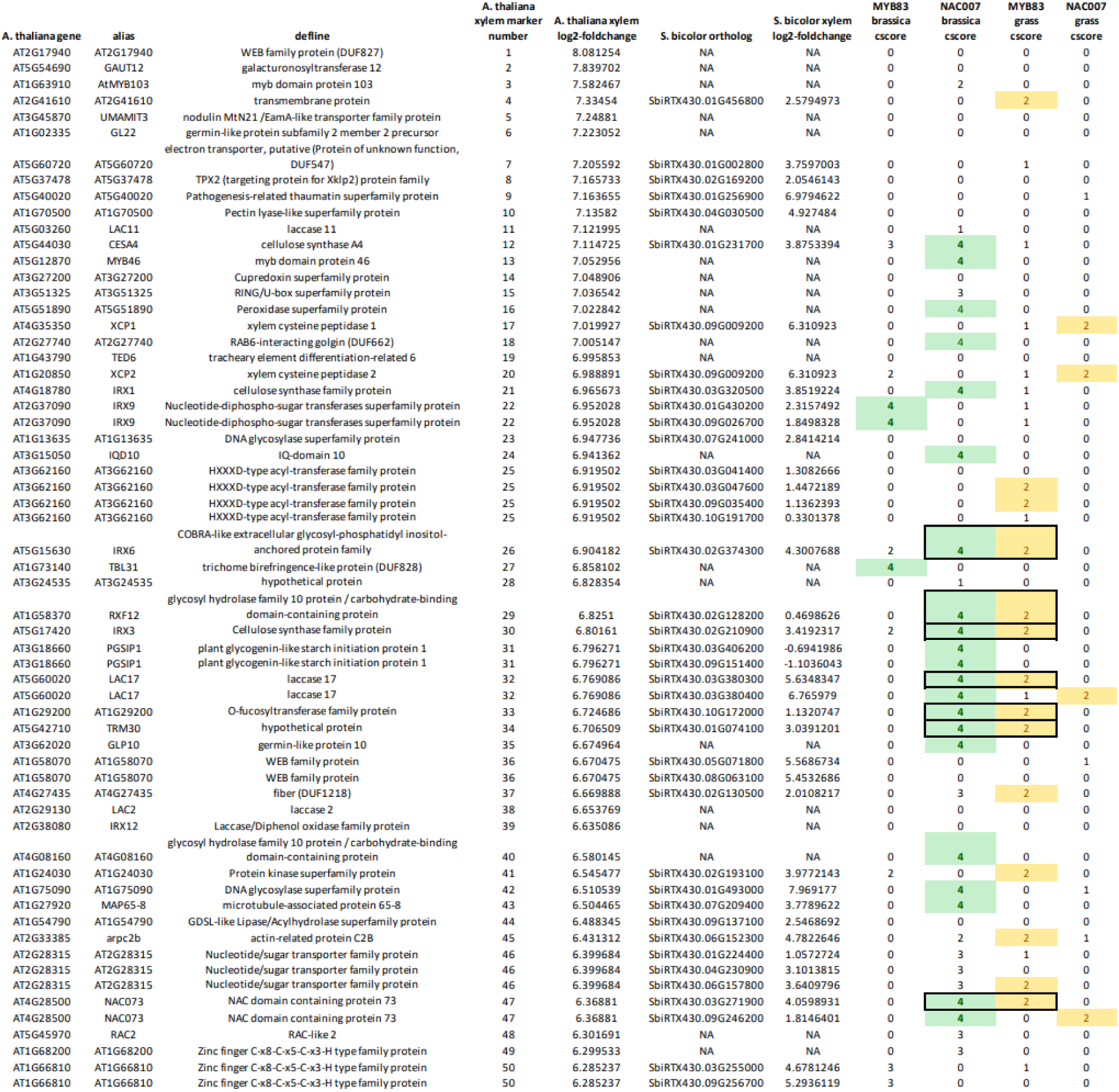
Table showing NAC007 and MYB83 TFBSs associated with the top 50 marker genes for the mature-xylem cluster in the *A. thaliana* seedling atlas and their sorghum orthologs. Specificity of expression of each ortholog in the mature xylem cluster of the sorghum root atlas is also shown in the column after the sorghum ortholog ID. Green shading highlights brassica-c4 TFBSs and yellow highlights grass-c2 TFBSs. Outlined boxes highlight *A. thaliana* genes with a brassica-specific NAC007 TFBS and no MYB83 TFBS whose sorghum orthologs have the reverse: a grass-specific MYB83 TFBS but no NAC007 TFBS.

